# S100A8-enriched microglia populate the brain of tau-seeded and accelerated aging mice

**DOI:** 10.1101/2023.11.10.566543

**Authors:** Roxane Gruel, Baukje Bijnens, Johanna Van Den Daele, Sofie Thys, Roland Willems, Dirk Wuyts, Debby Van Dam, Peter Verstraelen, Rosanne Verboven, Jana Roels, Niels Vandamme, Renzo Mancuso, Juan Diego Pita-Almenar, Winnok H. De Vos

## Abstract

Long considered to fluctuate between pro- and anti-inflammatory states, it has now become evident that microglia occupy a variegated phenotypic landscape with relevance to aging and neurodegeneration. However, whether specific microglial subsets converge in or contribute to both processes that eventually affect brain function is less clear. To investigate this, we analyzed microglial heterogeneity in a tauopathy mouse model (K18-seeded P301L) and an accelerated aging model (senescence accelerated mouse prone 8, SAMP8) using cellular indexing of transcriptomes and epitopes by sequencing. We found that widespread tau pathology in K18-seeded P301L mice caused a significant change in the number and morphology of microglia, but only a mild overrepresentation of disease-associated microglia. At the cell population-level, we observed a marked upregulation of the calprotectin-encoding genes *S100a8* and *S100a9*. In 9-months-old SAMP8 mice, we identified a unique microglial subpopulation that showed partial similarity with the disease-associated microglia phenotype and was additionally characterized by a high expression of the same calprotectin gene set. Immunostaining for S100A8 revealed that this population was enriched in the hippocampus, correlating with the cognitive impairment observed in this model. However, incomplete colocalization between their residence and markers of neuronal loss suggests regional specificity. Importantly, S100A8-positive microglia were also retrieved in brain biopsies of human AD and tauopathy patients as well as in a biopsy of an aged individual without reported pathology. Thus, the emergence of S100A8-positive microglia portrays a conspicuous commonality between accelerated aging and tauopathy progression, which may have relevance for ensuing brain dysfunction.

**Graphical abstract:** 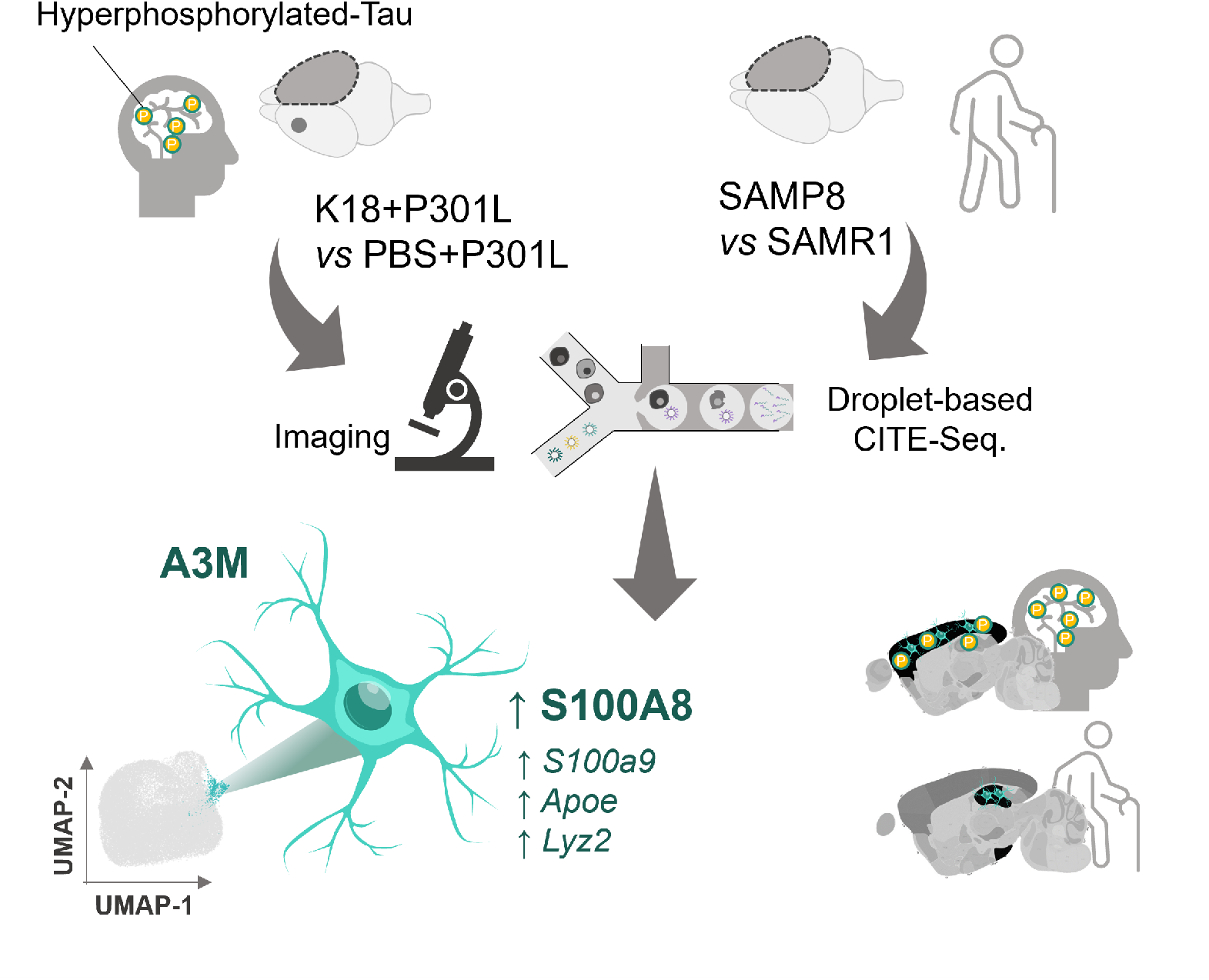

## Introduction

Microglia help maintain brain homeostasis by removing cellular debris and misfolded proteins and by pruning synapses to prevent excitotoxicity. A conspicuous property of microglia is their plasticity. Under normal physiological conditions, microglia display a ramified phenotype with compact cell body and elongated branching processes that probe the local microenvironment. When confronted with infection or tissue damage, they rapidly adopt an amoeboid morphology and migrate to the site of insult, where they upregulate their phagocytic capacity and secrete inflammatory cytokines and chemokines, such as Il-1ß (Hanisch & Kettenmann, 2007). This phenotypic switch is referred to as microglia “activation” and it is also observed in neurodegenerative disorders such as Alzheimer’s disease (Hanisch & Kettenmann, 2007). However, its contribution to brain dysfunction is not fully understood. For example, beneficial functions of activated microglia at early disease stages may have adverse effects when they persist in time, and hypofunctional microglia may promote chronic pathological conditions as well (Brelstaff et al., 2021). Indeed, by clearing protein deposits (e.g., amyloid ß) and damaged neurons (e.g., neurofibrillary tangle-bearing neurons), microglia may exert a neuroprotective function, but their abnormal or sustained activation can result in an exaggerated inflammatory response that triggers neuronal demise and promotes tau aggregation (Bhaskar et al., 2010; Gratuze et al., 2021; Mancuso et al., 2019). This points to a delicate balance in microglial functions, which is reflected by a growing spectrum of distinct phenotypes. Single-Cell RNA Sequencing (scRNA-Seq) approaches have revealed a core transcriptomic signature that is well-preserved among homeostatic microglia of the adult mouse (Li et al., 2019), and identified specific microglial subpopulations in neurodegenerative conditions, such as the disease-associated microglia (DAM) (Keren-Shaul et al., 2017). Most of this work has been done on mouse models that display amyloidosis, but since cognitive impairment correlates more strongly with tau deposition than with amyloid plaque load, and since a variety of other neurodegenerative disorders are solely caused by tau pathology, there is growing interest in tau as possible mediator of disease (Giannakopoulos et al., 2003). Recent studies suggest that microglial activation can accelerate tau aggregation and behavioural abnormalities in a hTau mouse model (Bhaskar et al., 2010; Chen et al., 2023; Gratuze et al., 2023; Sierksma et al., 2020; C. Wang et al., 2022), and that the accumulation of senescent microglia contributes to tau aggregation in a hTau.P301S (PS19) mouse model (Bussian et al., 2018). While there have been some studies investigating microglial subpopulations in models of tauopathy and in human brains (Chen et al., 2023; Gerrits et al., 2021; Kim et al., 2022), there is a lack of a comprehensive understanding of the influence of tau on microglia and the evolution of microglial heterogeneity during tau pathology progression. Furthermore, considering that microglia are long-lived cells, and that the most important risk factor for neurodegeneration is age, it can be expected that age-associated changes in microglia prime for pathology as well (Yoo & Kwon, 2022). Indeed, characteristic features such as a prolonged inflammatory response and lower phagocytic capacity may directly affect the accrual of pathogenic proteins. However, it is not known whether such features are confined to specific subpopulations and whether they occur in similar cells during aging and neurodegeneration. To better understand the evolution of microglial diversity upon tau pathology and aging, we here used Cellular Indexing of Transcriptomes and Epitopes by Sequencing (CITE-Seq) (Stoeckius et al., 2017) on two complementary mouse models and their respective controls, namely a K18-seeded (vs. PBS-injected) P301L mouse model displaying progressive tau pathology and a Senescence Accelerated Mouse Prone 8 (SAMP8, vs. Senescence-Accelerated-Resistant Mouse, SAMR1) mouse model showing accelerated aging. In doing so, we identified specific microglia in both models that express high levels of the calprotectin encoding genes *S100a8* and *S100a9* and which represent a distinct accelerated aging-associated sub-population in older SAMP8 mice.

## Materials and methods

### Mice handling

In this study, 206 mice (*n*) were used: C57Bl6/J mice (*n* = 48), Heterozygote CX3CR-1^+/GFP^ mice (*n* = 9), SAMP8/TaHsd mice (*n* = 28), SAMR1/TaHsd mice (*n* = 30), FVB/NJ mice (*n* = 28) and hTau.P301L (*n* = 66) (Supplementary Table 1) (Jung et al., 2000; Terwel et al., 2005; Yagi, Katoh, Akiguchi, & Takeda, 1988). The mice were respectively named C57Bl6/J, CX3CR1^+/GFP^, SAMP8, SAMR1, FVB and P301L in the article.

All mice were housed under a 12 h light/dark cycle, with food and water supplied *ad libitum* and with cage enrichment. For PLX3397 and stereotaxic injections, mice were single housed and randomised per treated group, considering multiple assignments within the same litter. The experiment and animal environment were carefully adjusted to minimize confounding factors. The investigators responsible for performing the surgeries, administering treatments, and conducting behavioural tests were blinded to the experimental conditions.

All experiments were performed in accordance with the EU Directive 2010/63/EU and protocols approved under ECD files 2020-65 (University of Antwerp) and ECD protocol 628_Aggregate spread (Janssen Pharmaceutica). Methods and results are written in accordance with the ARRIVE guidelines 2.0 for publishing in vivo research.

### Stereotactic injections

K18 and PBS injections were done as previously described (Detrez et al., 2019), with slight modifications, at Janssen Laboratory (Beerse, Belgium). Briefly, at the age of 90 ± 5 days, FVB and P301L mice were deeply anaesthetised with isoflurane (2% in 36% oxygen) and fixed in a stereotactic frame (Neurostar, Germany, Stereodrive software v 2019). A 30G syringe (Hamilton) was used for injecting 2 μl of PBS or K18 (7.5 µg) in the right hemisphere at a speed of 0.2 μl/min at the selected coordinates: anterior-posterior −2.0 mm, medial-lateral +1.6 mm from Bregma, and dorsal-ventral +1.4 mm from the dura.

### Microglia depletion

Mice were fed at Janssen Laboratory (Beerse, Belgium) *ad libitum* using modified AIN76A (15.5% Dextrose, 4% sucrose) chow ((Bio-Services BV) supplemented with the Colony Stimulating Factor 1 Receptor (CSF1R) inhibitor PLX3397 (290 mg of PLX3397/kg chow) or PLX5622 (1200 mg of PLX5622/kg chow). The 3-months-old CX3CR1^+/GFP^ mice were treated during 7, 14, 21 or 28 days. For FVB and P301L mice, the PLX3397 treatment began 14 days before stereotactic injection and was sustained until 3 months post-injection.

### Perfusion and brain extraction

The animals designated for CITE-Seq were transported to the Center for Inflammation Research at VIB-UGent (Ghent, Belgium) the day before the experiment. On the day of the experiment, they were deeply anesthetized by intraperitoneal injection (Nembutal, 150 mg/kg), followed by transcardial perfusion with ice-cold heparinized PBS (Sigma H3393-50KU; 10 U/ml; 4 ml/min). Thereafter, brains were extracted, hemisected and stripped of the olfactory bulb and cerebellum for subsequent cell dissociation.

For FVB and P301L brain microscopy, animals were sequentially transcardially perfused with PBS at Janssen Laboratory (Beerse, Belgium) 4% paraformaldehyde (PFA) (Affymetrix USB. J19943; 5 min) at 4 ml/min. Thereafter, brains were hemisected and post-fixed overnight in 4% PFA at 4 °C, followed by a PBS wash (3 × 15 min) and stored in PBS with 0.1% NaN3 at 4 °C until further processing.

For microscopy analysis of SAMP8 and SAMR1 mice, the animals were perfused sequentially with PBS at the Laboratory of Cell Biology & Histology (Wilrijk, Belgium) and post-fixed overnight in 4% PFA at 4 °C. Afterward, they underwent PBS washing and were stored in PBS with 0.1% NaN3 at 4 °C until further processing.

### Flow cytometry and flow-assisted cell sorting

Contralateral hemispheres (*i.e.,* opposite to the injected hemisphere) were subjected to mechanical and enzymatic dissociation using the GentleMACS™ dissociator (130-093-235, Miltenyi Biotec) in combination with the Neural Tissue Dissociation Kit (130-092-628, Miltenyi Biotec), according to the manufacturer’s instructions. To avoid microglial activation during extraction, we dissociated the hemispheres in the presence of Actinomycin D (5 nM, Sigma-Aldrich SBR00013) and Brefeldin A (1×, 420601 BioLegend) (Supplementary Fig. 1). After enzymatic dissociation, cells were resuspended in 30% Percoll (P1644, Sigma-Aldrich) and centrifuged for 15 min at 300 g at 4°C for myelin removal. The supernatant containing the myelin was removed, and the pelleted cells were washed with FACS buffer (PBS, 0.5 % BSA, 2 mM EDTA). Up to 2 million cells per mouse were incubated for 30 min on ice with 25µL of staining mix in PBS containing 0.04% BSA, Cluster of Differentiation CD11b-PE-Cyanine5 antibody (BioLegend, cat 101210), TruStain FcX Block (BioLegend, cat 101320) and a mouse CITE-Seq antibody panel. The CITE-Seq pane contains 158 unique oligo-conjugated antibodies and isotype controls (TotalSeq™-A) (Supplementary Table 2). Multiple replicates were tagged and pooled using TotalSeq-A hashtag antibodies.

### Cellular Indexing of Transcriptomes and Epitopes by Sequencing

FACS Aria II-sorted single-cell suspensions were resuspended at an estimated final concentration of 1000 cells/µl and loaded on a Chromium GemCode Single Cell Instrument (10x Genomics) to generate single-cell Gel beads-in-EMulsion (GEM). The DNA libraries were prepared using the GemCode Single Cell 3’ Gel Bead and Library kit, version NextGEM 3.1 (10x Genomics) according to the manufacturer’s instructions with the addition of amplification primers (3nM, 5’CCTTGGCACCCGAGAATT*C*C and 5’GTGACTGGAGTTCAGACGTGTGC*T*C) during cDNA amplification to enrich the TotalSeq-A cell surface protein and hashtag oligos. Size selection with SPRIselect Reagent Kit (Beckman Coulter, B23318) was used to separate amplified cDNA molecules for 3’ gene expression and cell surface protein construction. TotalSeq-A protein library construction including sample index PCR using Illumina’s Truseq Small RNA primer sets and SPRIselect size selection was performed according to the manufacturer’s instructions. The cDNA content of pre-fragmentation and post-sample index PCR samples was analyzed using the 2100 BioAnalyzer (Agilent). Sequencing libraries were loaded on an Illumina NovaSeq flow cell at VIB Nucleomics core with sequencing settings (Supplementary Table 3) according to the recommendations of 10x Genomics, pooled in an 80:25 ratio for the combined 3’ gene expression and cell surface protein samples, respectively. The Cell Ranger pipeline (10x Genomics, version 6.0.0) was used to perform sample demultiplexing and to generate FASTQ files for read 1, read 2 and the i7 sample index for the gene expression and cell surface protein libraries. Read 2 of the gene expression libraries was mapped to the reference genome (mouse mm10) using STAR. Subsequent barcode processing, unique molecular identifiers filtering, and gene counting was performed using the Cell Ranger suite version 6.0.0 (10X Genomics) and Seurat v4.0.5. For further analysis, we used the filtered feature-barcode matrix generated by CellRanger (v6.0.0, 10X Genomics).

### Data processing and quality control

Data processing was performed in R 4.1.2. Individual libraries were processed in Seurat (v4.0.5) (Hao et al., 2021). Libraries were filtered excluding cells with less than three features and genes detected and features and genes present in less than 200 cells. All three modalities (Gene EXpression (GEX), HashTag Oligonucleotide (HTO) and Antibody-Derived Tag (ADT)) were normalized. Individual replicates were extracted with Seurat’s HTODemux, and doublets were excluded per library. Individual libraries were filtered based on the number of UMI’s lower than mean + 2sd, number of genes between mean ± 2sd and percent mitochondrial DNA lower than mean + 2sd. Before integration 2000 variable features were detected per library. Libraries were integrated with Seurat’s Integrate Data. Initially, cells clustered into distinct populations based on the number of UMI’s, the number of genes and the mitochondrial DNA, so these variables were regressed out with ScaleData. Non-microglial cells were excluded from the final objects, including peripheral macrophage (*Mrc1, Cd163*), neutrophils (*Retnlg, Mrsb1*), macrophages (*Plac8, Adgre5*), proliferating cells (*Top2a, Mki67*), choroid plexus cells (*Ttr*), endothelial cells (*Cldn5*, *Ly6c1*) and doublets/low quality cells (*Atp1a2*, *S100a9*) and other immune cells (*Gzma*, *Nkg7*). A principal component analysis (PCA) was used to reduce the dimensionality of the dataset. Uniform Manifold Approximation and Projection (UMAP) space and neighbourhood embeddings were calculated based on 18 PC’s. Finally, clustering was performed at a resolution of 0.4 resulting in six distinct microglia subpopulations mice for SAMR1 and SAMP8 mice and at a resolution of 0.6 resulting in five distinct microglia subpopulations for injected P301L and FVB mice. Differential expression testing was performed using Seurat’s FindMarkers. Genes with an adjusted p-value of <0.05 and absolute log2FC of > ±0.25 were considered as differentially expressed. All plots were generated in R with the ggplot2 package (version 3.3.5) or EnhancedVolcano (version 1.10.0). The top 10 representative genes or proteins of each cluster were depicting using dot plot. For each dot plot, the colour of the dots represents the normalized average of expression of the gene in the cluster and the size of the dot represents the percentage of cells expressing this gene. Volcano plots were used to depict differential expression results between groups. Only significant genes after wilcox test (P_value adjusted < 0.05) that show fold-change > 0.25 are represented. Genes that present in addition an absolute fold-change > 1 or 0.5 are represented in red.

To investigate enrichment of the ageing markers in the P301L dataset, distinctive genes for the accelerated ageing signature were defined by FindMarkers in Seurat of the ageing associated microglia versus disease associated microglia in the SAMP8 object. From here the top 30 markers were extracted. Enrichment scores were calculated with AddModuleScore in Seurat.

### Pseudotime analysis

Pseudotime analysis was performed with Monocle3 (v1.0.0) (Almanzar et al., 2020; McInnes, Healy, & Melville, 2018; Trapnell et al., 2014). The Seurat object was converted into a CellDataObject with SeuratWrappers (v 0.3.0). The object was preprocessed with 100 dimensions and cells were clustered. The cells were ordered with order_cells and the cluster of homeostatic microglia as the starting point. Two trajectories were extracted from the SAMR1/SAMP8 object with choose_graph_segments; one from homeostatic microglia to DAM and one from the homeostatic microglia to accelerated aging-associated microglia. For both branches models were fitted with “∼model” before running differential expression testing across the branch, resulting in a dataset with genes that are upregulated across pseudotime.

### Immunofluorescence staining

After perfusion and PFA 4% fixation, fresh tissue was dissected into coronal 30 µm sections using a vibratome (Leica) and collected in PBS with 0.1% NaN3 or cryopreserved brains were sliced into sagittal 12 µm sections using a cryostat (Leica) for pentameric formyl thiophene acetic acid (pFTAA) staining. After epitope retrieval using sodium citrate buffer (10 mM, pH 6.0) at 95°C during 10 min (required for S100A8 staining) and PBS wash, the sections were incubated with a blocking buffer containing 0.3% Triton X-100 and 5% normal horse serum diluted in PBS for 1h30 at room temperature (RT). Subsequently, slices were incubated with primary antibodies diluted in blocking buffer overnight at 4°C or 2h at RT for pFTAA (3 µM, gift from Dr. Peter Nilsson). The primary antibodies used were NeuN (1:1000, guinea pig (GP), ABN90P, Millipore), SOX9 (1:200, rabbit, 82630s, Cell Signalling Technology), SOX10 (1:500, goat, AF2864, R&D systems), Ionized calcium Binding Adaptor molecule 1 (IBA1) (1:500, rabbit, 019-19741, Wako), CD63 (1:200, rat, 143902, Biolegend), S100A8/MRP8 (1:250, rabbit, ab92331, Abcam), IBA1 (1:500, goat, ab5076, Abcam), S100A9 (1:200, rat, ab105472, Abcam), P2RY12 (1:100, rat, 848002, Biolegend), CD11b (1:400, rabbit, LS-C141892, lifespan biosciences), Ly6G (1:200, rat, BE0075-1, BioCell). After a PBS wash, sections were incubated with donkey secondary antibodies (Jackson Immuno Research) labelled with FITC, Cyanine-3 or Cyanine-5 and diluted (1:500) in blocking buffer. After several washes, slices were incubated for 5 min with 5µg/ml of DAPI and were mounted on slides using Citifluor^TM^ Mounting solution AF-1. Confocal images were acquired of the complete slices (as tile scan) with a PerkinElmer UltraVIEW Vox spinning disk confocal system using a 20x/NA 0.5 or 10x/NA 0.35 objective (yielding a pixel size of 0.36 × 0.36 µm^2^, resp. 0.72 × 0.72 µm^2^), or with a Nikon Ti2 W1 spinning disk confocal using a 20x/NA 0.75 or 10x/NA 0.45 objective (yielding a pixel size of 0.325 × 0.325 µm^2^, resp. 0.65 × 0.65 µm²).

### Human brain tissue

Brain sections from the transentorhinal region designated for immunostaining were obtained from the NeuroBiobank of the Institute Born-Bunge (NBB-IBB), Wilrijk (Antwerp), Belgium (ID: BB190113) and donors gave informed consent to donate their brain to the NBB-IBB. Ethical approval was granted by the ethics committee of the University Hospital of Antwerp and the University of Antwerp (Antwerp, Belgium) (project ID 5880). The cohort comprised of 28 individuals ranging from 40 to 70 years, with an equal distribution of female and male subjects. This included 10 patients diagnosed with Alzheimer’s Disease (AD) at a Montine pathology stage of A3B3C3, 8 patients with various forms of Tauopathy, and 10 controls with no significant neuropathological findings (Supplementary Table 4).

The right brain hemisphere was fixed in buffered 10% formaldehyde, neutral pH for 6 to 12 weeks, dissected by brain region into embedding cassettes and processed into formalin-fixed, paraffin-embedded slides (FFPE). FFPE slides of 5 micrometres were prepared with a microtome (Thermo Fisher Scientific, Microm HM 355S).

Human paraffin-embedded tissue sections underwent de-paraffinization using xylene and ethanol (100% to 80%). Antigen retrieval was performed for 40 minutes at 95°C in sodium citrate buffer (10 mM, pH 6.0). After washing with PBS, sections were permeabilized for 2 hours at RT in PBS containing 0.05% Tween20 and subsequently further permeabilized and blocked using a blocking buffer consisting of 1% Triton X-100 and 5% normal horse serum in PBS at RT. The slices were stained with S100A8/MRP8 (1:250, rabbit, ab92331, Abcam) and IBA1 (1:500, goat, ab5076, Abcam), mounted, and acquired using a Nikon Ti2 W1 spinning disk confocal with a 10x/NA 0.45 objective, as described previously.

### Image analysis

Confocal tiles were stitched using Volocity (PerkinElmer) or NIS elements (Nikon), Z-stacks were flattened by a maximum projection and background corrected in ImageJ (Schneider, Rasband, & Eliceiri, 2012) and quantifications were performed in QuPath (Bankhead et al., 2017). Brain regions were first manually delineated and annotated. Nucleus detection and segmentation was done using the CellDetection plugin or StarDist (Schmidt, Weigert, Broaddus, & Myers, 2018), after which the signal intensity in the respective antibody channels was measured per nucleus. The number of positive cells for a given marker was calculated using R studio (version 2022.7.1.554) using a fixed threshold on the average nuclear intensity per marker. The number of positive cells was normalized to the total area of the brain slice. Regarding the quantification of S100A8^+^IBA1^+^ cells and IBA1^+^CD63^+^ cells, a semi-blinded approach was employed by the investigator. Double-positive cells were manually counted and then normalized to the total number of IBA1^+^ positive cells, which were determined as described above.

### Whole hemisphere imaging and analysis

Fluorescent labelling and clearing of brain hemispheres were done based on the Immunolabeling-enabled three-Dimensional Imaging of Solvent-Cleared Organs plus (iDISCO+) as done before (Detrez et al., 2019). Briefly, after dehydration, autofluorescence reduction and rehydration using different methanol baths, brains were incubated during 2 weeks at 37°C in presence of an AT8 antibody that specifically binds hyperphosphorylated tau (pSer202/Thr205/PSer208, produced at Janssen Pharmaceutica) and is directly labelled with a near-infrared fluorescent tag (PerkinElmer VivoTag 680XL). Brain samples were then dehydrated, cleared and acquired with an Ultramicroscope II (Lavision Biotec GmbH), equipped with an Olympus MVPLAPO 2× (NA 0.50) objective lens and DBE-corrected LV OM DCC20 dipping cap. Images were recorded with a Neo sCMOS camera (Andor) with a magnification of 1.6x/NA 0.5 and 10 µm axial sampling, resulting in 2 × 2 × 10 μm^3^ voxels. Images from the left and right light sheets were merged on the fly with a linear blending algorithm. A 488 nm (for autofluorescence) and 640 nm (for AT8) laser with a 525/50 nm and 680/ 30 nm emission filter were used. Sagittal optical sections were recorded in a mosaic of two tiles. Automated analysis of whole-hemisphere microscopy images was performed using a previously described pipeline (Detrez et al., 2019). Briefly, the autofluorescence channel was aligned to a 3D light sheet reference brain atlas, and the resulting transformation vector set was used for regional analysis of the AT8 signal. The total number of detected voxels for a given brain region was calculated and normalized to the volume of the respective region.

### Statistical analysis

The determination of the required number of mice for whole hemisphere imaging, immunofluorescence staining on slices, and CITE-Seq was based on previous studies (Detrez et al., 2019) and guided by the use of Statulator, an online statistical calculator (With a permitted type I error (*α*) of 0.05 and power (1 − *β*) of 0.80). Values are reported as mean ± standard error of the mean (SEM). Normality of data was assessed using the Kolmogorov– Smirnov test. For a comparison between two conditions, a parametric one-tailed t-test or a non-parametric one-tailed Mann-Whitney U test was performed, depending on the normality checks. For comparison between three or more conditions, one-way or two-way ANOVA with Dunn’s, Tukey’s or Sidak’s multiple comparisons test was performed. For CITE-Seq, Seurat’s FindMarkers was used in combination of Wilcox test. Statistical analyses were performed as indicated in the figure legends using GraphPad Prism (GraphPad Software Inc. version 9.0.1) or R studio (version 2022.7.1.554) for CITE-Seq data.

### Exclusion criteria

Animals were excluded from the study or euthanized if any of the following conditions occured: meeting any defined humane endpoint criteria, having open, bleeding, or infected wounds that cannot be healed due to aggressive behaviour from cage mates, or experiencing sudden death in the cage. Outliers identified using GraphPad Prism and determined by ROUT test (with a *Q* value of 1%) were excluded from the analysis. The K18+P301L mice processed for CITE-seq analysis at 1-week post-injection were excluded from the study due to a low cell count.

## Results

### Microglia depletion reduces regional tau load in K18-seeded P301L mice

To shed light on the contribution of microglia to tau pathology development, we first quantified the impact of their depletion in a K18-seeded P301L mouse model (P301L+K18) on hyperphosphorylated tau spreading. P301L mice are transgenic FVB mice engineered to express a specific mutant form of the human tau protein known as the 4R/2N isoform. They display slow endogenous accrual of hyperphosphorylated tau, with visible tau pathology after 7 months of age, primarily in the cerebellum and brainstem. Stereotactic injection with K18, a truncated form of human tau containing only the 4 microtubule binding repeats, in the CA1 region at 3 months of age, drastically accelerates this process (Peeraer et al., 2015). Importantly, it more faithfully reproduces the characteristic tau spreading pattern observed in human Alzheimer’s disease patients compared to non-seeded P301L mice (Detrez et al., 2019). For microglial depletion, we selected the CSF1R inhibitor PLX3397 (over PLX5622) as in our hands, it consistently reduced the number of microglia to less than 5% of the endogenous population without affecting the other cell types in a CX3CR1^+/GFP^ mouse model (Supplementary Fig. 2). When fed with PLX3397-supplemented chow for 104 days (14 days pre-+ 3 months post-injection, p.i.), P301L mice showed a sustained depletion of IBA1-positive microglia as well, despite their higher baseline levels than FVB control mice (Supplementary Fig. 3). We then quantified tau pathology in whole brain after AT8 immunostaining and iDISCO+ clearing. In accordance with our previous findings (Detrez et al., 2020, 2019), buffer-injected transgenic (P301L+PBS) and non-transgenic (FVB+PBS) animals did not show significant AT8 staining at this stage, whereas age-matched K18-injected P301L mice showed widespread AT8-positivity (Fig. 1A, Supplementary Fig. 4). When comparing the total AT8-positive pixel fraction (further referred to as phospho-tau load) between P301L+K18 mice fed with standard or PLX3397-supplemented chow, we found a brain-region dependent decrease in the latter. While the phospho-tau load in the ipsilateral hemisphere remained equally high after PLX3397 treatment (Supplementary Fig. 4), it was significantly reduced in specific cerebral regions (incl. the isocortex (*P* = 0.0003), Ammon’s horn of the hippocampus (*P* = 0.0411) and striatal dorsal region (*P* = 0.0175)) of the contralateral hemisphere (Fig. 1B). Visual inspection of contralateral K18-seeded P301L mouse brain sections revealed, next to the marked upregulation of IBA1 signal, conspicuous morphological changes of microglia (from ramified to rod-like or amoeboid), especially in the hippocampus, in line with our previous observations (Detrez et al., 2019). However, other regions that were affected by PLX3397 treatment, such as the somatomotor area in the isocortex, did not reveal a distinct switch in microglial phenotype (Fig. 1C). Thus, we conclude that PLX3397 treatment mitigates tau spreading in K18-seeded P301L mice but that there is no one-to-one relationship with the microglial morphotype.

**Figure 1.**
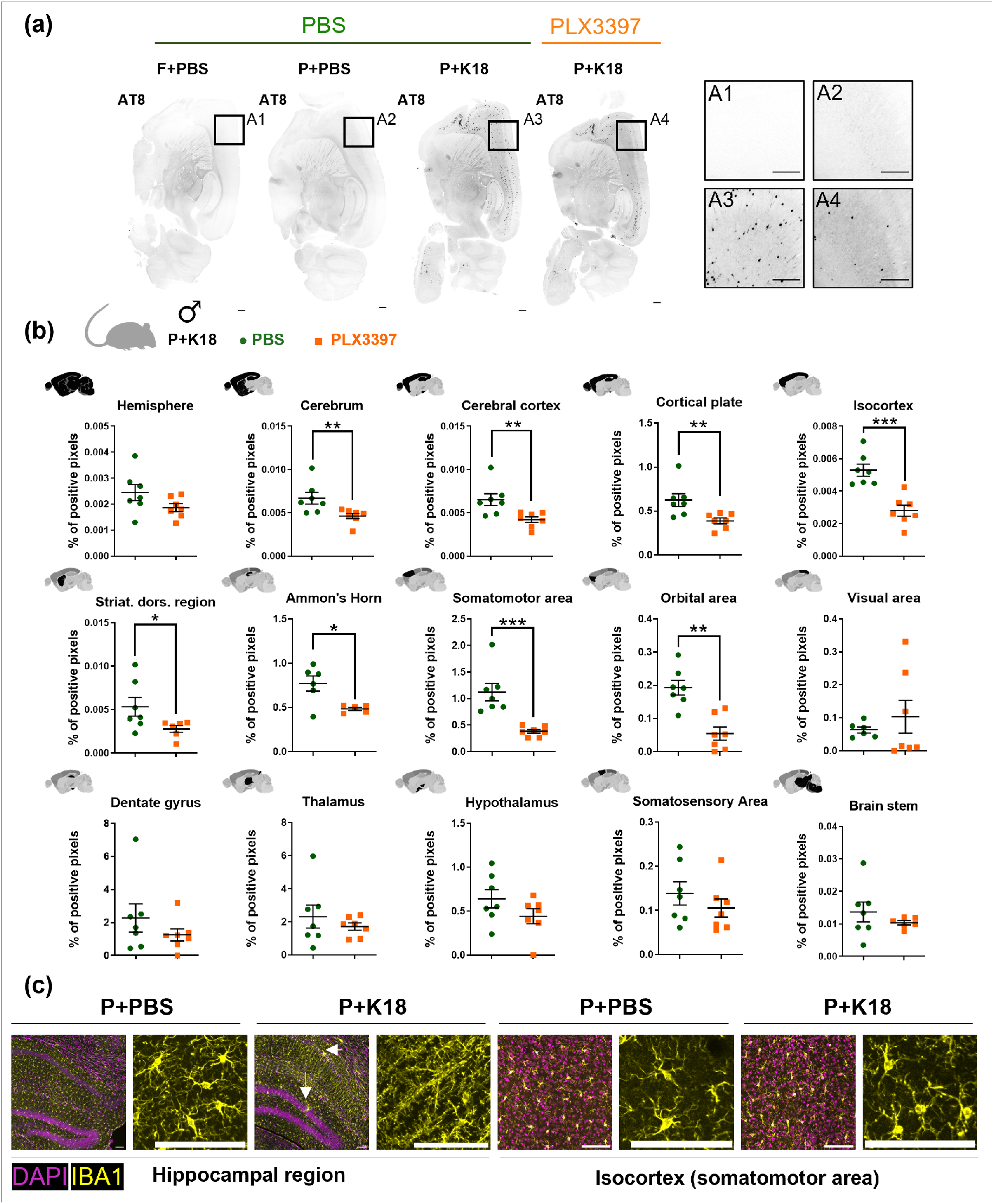
Microglia depletion reduces regional tau load in contralateral hemisphere of K18-seeded P301L mice. (**a**) Representative contrast-matched images of AT8 staining of sagittal slices of the contralateral hemisphere of PLX3397 treated K18-injected P301L (P+K18) and non-treated PBS-injected FVB (F+PBS), PBS-injected (P+PBS) and K18-injected P301L (P+K18) male mice. Insets represent zoomed-in views on the isocortex. Scale bar 400 µm. (**b**) Quantification of AT8 staining in different regions of the contralateral hemisphere of P+K18 after 104 days (14 days pre-+ 3 months post-injection) of feeding with modified AIN76A supplemented with PBS (green dot) or PLX3397 (orange square). Positive pixels were quantified and normalized over the total number of pixels (% of positive pixels) in different regions (represented in black). (**c**) Representative images illustrating IBA1 and DAPI staining in the hippocampal region and the isocortex (somatomotor area) of both P+PBS and P+K18 mice, with zoomed-in sections to highlight microglial morphology. White arrows indicate ameboid morphology, which is not depicted in the zoomed image. Scale bar 100 µm. Values are mean ± SEM. Outliers were detected using ROUT test (*Q* = 1 %) and excluded from analysis. Statistical differences (\**P* < 0.05, \*\**P* < 0.01 and \*\*\**P* < 0.001) were determined by non-parametric one-tailed Mann-Whitney U test or parametric one-tailed t-test depending on normality checks. (*n* = 7 mice/condition) Striat. dors. region = striatum dorsal region

### Distinct microglial phenotypes populate P301L mouse brain

Having established that K18 seeding triggers distinct microglial changes and that PLX3397 treatment affects tau pathology, we next asked whether this would align with the selective enrichment of specific microglial subpopulations in this model. Hence, we mapped the microglial heterogeneity in the contralateral hemispheres of K18-seeded P301L mice and their buffer-injected controls (P301L+PBS and FVB+PBS) at 1 week, 1 month and 3 months p.i. using CITE-Seq after CD11b-based FACS enrichment (Fig. 2A). We will further refer to this dataset as the P301L dataset. After PCA-based dimensionality reduction and graph-based clustering, we found that most of the CD11b-positive cell population consisted of non-proliferating microglia (87 %). Other cell types including peripheral macrophages (10%), macrophages (1%), neutrophils (1%), proliferating microglia (1%), and choroid plexus cells (0.20 %) were excluded from further analysis (Fig. 2B; Supplementary Fig. 5). After exclusion, we retained 51,163 non-proliferating microglia for unsupervised clustering and represented them in UMAP space. This revealed the presence of five microglial subpopulations, which we identified based on their proteomic (epitopes) and gene expression (transcriptome) patterns into homeostatic microglia, ribosomal response microglia, early activation response microglia, DAM and Interferon Response Microglia (IRM) (Fig. 2C-F). Homeostatic microglia were the most abundant subpopulation (80%) and were characterized by their high level of homeostasis-associated transcripts (*e.g., Cx3cr1*, *Tmem119*, *Siglec*, *Gpr34*, *P2ry12* and *P2ry13)* and all-over low level of membrane markers. Ribosomal response microglia (12.5%) were enriched in ribosomal genes and presented high levels of integrin and receptors involved in cell adhesion and migration (*e.g., Cx3cr1*, *Cd54* and *Cd49f*). Early activation response microglia (4%) expressed higher levels of immediate early genes (e.g., *Jun*, *Junb*, *Fos*, *Dusp1* and *Egr1)* and interferon response microglia (0.45%) were characterized by transcripts of the interferon pathway (*e.g., Ifit2* and *Ifit3*). Finally, DAM (3%) displayed a distinct set of genes associated with lipid and lipoprotein metabolism (*Apoe*, *Lpl*) and lysosomal function (*Lyz2*) and presented T-cell related membrane proteins such as CD86, CD11c and CD63.

**Figure 2.**
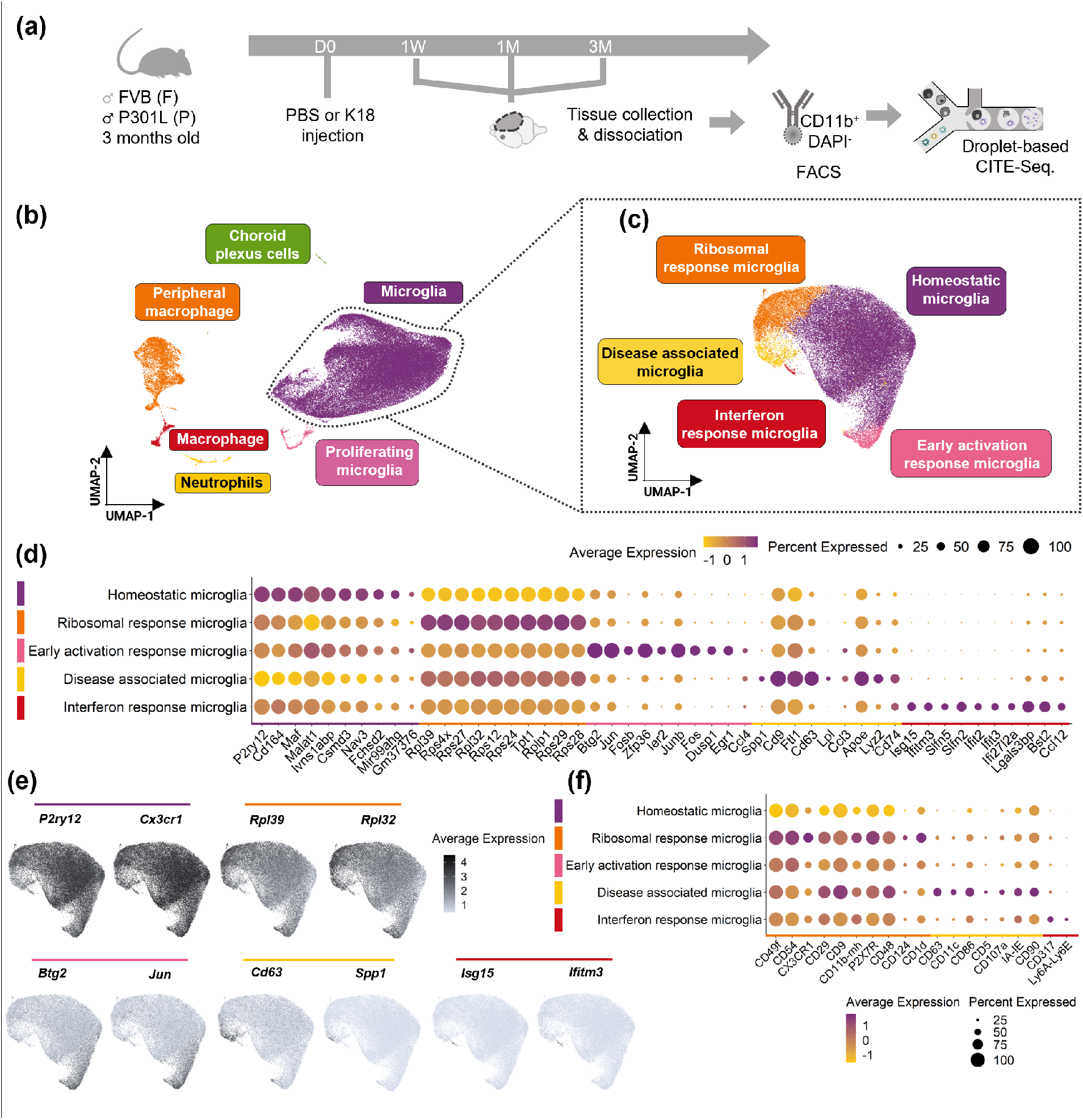
Characterization of microglial subpopulations in injected FVB and P301L mice. (**a**) Schematic diagram showing the CITE-Seq timeline. (**b**) UMAP plot of CD11b^+^ selected cells depicting the different cell types across all different groups (59129 cells). (**c**) UMAP plot of microglia depicting the different microglial subpopulations of all different groups (51163 cells). (**d**) Dot plot representing the top 10 differentially expressed genes of each microglial subpopulation. (**e**) Feature plot depicting the relative expression of two representative genes of each microglial cluster. (**f**) Dot plot displaying the representative protein (epitope) markers of each microglial subpopulation determined by CITE-sequencing. (*n* = 4 mice/condition).

### K18-seeding induces a mild increase of DAM in P301L mice

Upon deconvolution of the UMAP plot, we found that all five microglial subpopulations were present in each mouse model of the P301L dataset (P310L+K18, P301L+PBS, FVB+PBS) indicating that induced tau pathology does not give rise to (or loss of) a specific microglial subpopulation (Fig. 3A). However, when further refining the analysis to the individual time points (the 1-week time point was not analysed due to its lower cell number), we noticed that the DAM population was specifically elevated in the P301L+K18 model 3 months p.i. compared to its controls, but only significantly with respect to FVB+PBS (*P* = 0.0427) (Fig. 3B). Using CD63 as one of its most discriminatory membrane markers (Fig. 2F, Fig. 3C), we quantitatively confirmed this significant increase of the DAM population in brain sections (P301L+K18 *vs* FVB+PBS, *P* = 0.0477) (Fig. 3D-E). Herein, we found DAM to be predominantly located in the dorsal region of the brain, including the brainstem, corticospinal tract and the striatum. Despite a similar trend towards increase, no significant differences were observed at 1-month p.i. All-over, the frequency of DAM was low (<5%). Therefore, we conclude that K18 seeding induces a mild increase of DAM in P301L mice but otherwise leaves the global microglial composition roughly unaltered.

**Figure 3.**
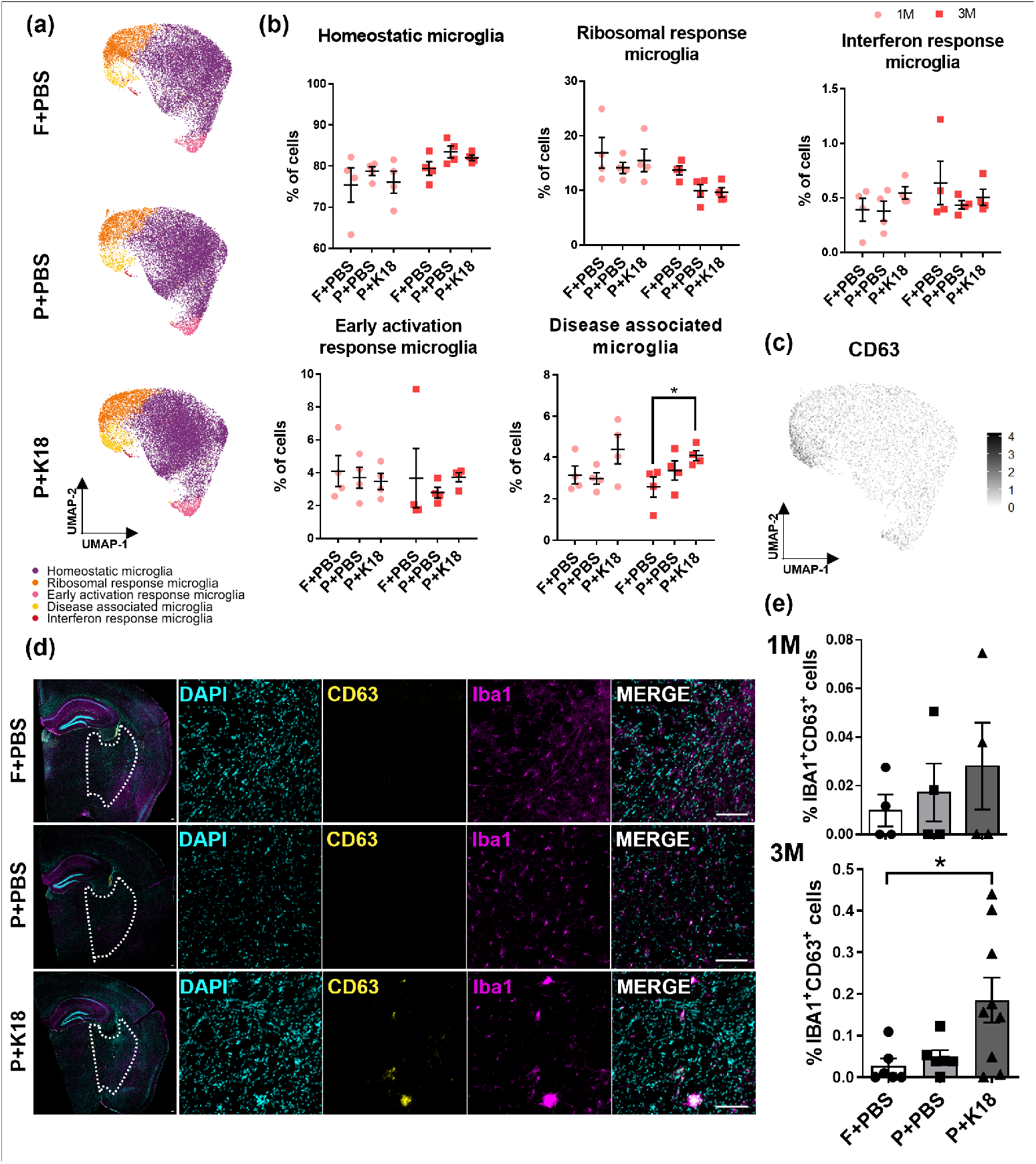
DAM subpopulation is increased in K18-seeded P301L mice at 3 months post-injection. (**a**) UMAP plots depicting the different microglial subpopulation in PBS-injected FVB (F+PBS), PBS-injected (P+PBS) and K18-injected (P+K18) P301L mice. (**b**) Quantification of the percentage of cells in each microglial subpopulation over the different models at 1-month and 3-months post-injection. (**c**) CD63 protein level projected onto the UMAP plot shows its enrichment in DAM. (**d**) Representative images at 3-months post-injection of CD63^+^IBA1^+^ cells. Images represent a merge of a whole hemisphere slice (left) and a zoom in the delineated region were IBA1^+^CD63^+^ cells were localized. (**e**) Quantification of the percentage of CD63^+^IBA1^+^ cells over the total number of IBA1^+^ cells in the delimited region (**d** dotted line) at 1- and 3-months post-injection in the different mice models (2-3 slices/mouse). Scale bar 100 µm. Values are mean ± SEM. Statistical difference (\**P* < 0.05) was determined by one-way ANOVA with Dunn’s multiple comparisons test for each timepoint. (*n* = 4-7 mice/condition)

### Microglia of older P301L+K18 mice express calprotectin-encoding genes

To scrutinize the global transcriptional changes within the microglial population, we next performed a Differential Gene Expression (DGE) analysis between buffer-injected and K18-seeded P301L mice. This revealed a limited subset of dysregulated genes with significant p-values (Fig. 4A), which mainly corresponded with those portraying ribosomal response (*Rpl* and *Rps* gene families) and DAM phenotypes (*Apoe* and *Lyz2*). However, at 3-months p.i., the most prominently upregulated genes were *S100a8* and *S100a9*, which encode calcium-binding cytosolic proteins that together form the pro-inflammatory complex calprotectin (S. Wang et al., 2018). The expression of these genes was not specific to any of the identified microglial sub-populations (Fig. 4B-C). Since both genes were not upregulated at 1-month p.i., these data suggest that they are a consequence and not an early mediator of the induced tau pathology.

**Figure 4.**
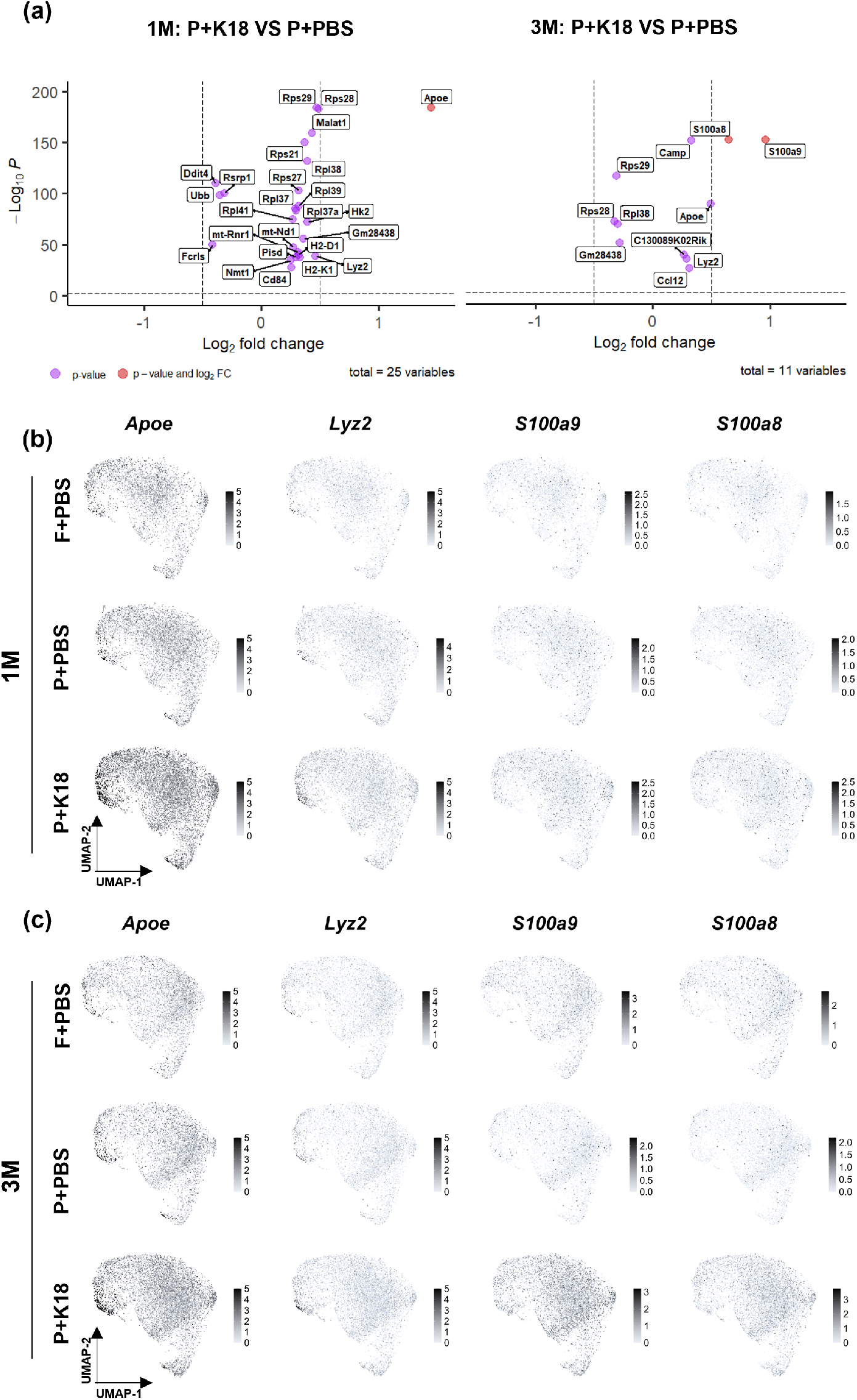
In K18-seeded P301L mice, microglia exhibit high expression levels of *S100a8*, *S100a9*, *Apoe* and *Lyz2*. (**a**) Volcano plot depicting the differentially expressed genes between PBS-(P+PBS) and K18-injected (P+K18) mice at 1-month and 3-months post-injection. (**b-c**) Feature plot depicting the relative level of expression of *Lyz2*, *Apoe*, *S100a8*, *S100a9* genes in the different group at 1-(**b**) and 3-(**c**) months post-injection.

### Microglia in SAMP8 accelerated aging mice display changes in metabolic and immune gene expression

*S100a8* and *S100a9* levels have been shown to increase with aging (Swindell et al., 2013), and emerging evidence suggests a link between hyperphosphorylated tau and accelerated aging (Ramirez et al., 2022). Furthermore, analysis of the CSF proteome in AD patients reveals accelerated biological aging of the innate immune system correlated with tau (Cullen, Mälarstig, Stomrud, Hansson, & Mattsson-Carlgren, 2021). Therefore, we hypothesized that the increased expression of *S100a8* and *S100a9* might reflect K18-induced accelerated brain aging. To investigate this, we analyzed using the same CITE-Seq procedure, the senescence accelerated mouse prone 8 (SAMP8) model, described to exhibit elevated tau levels and accelerated aging (Liu, Liu, & Shi, 2020), and its normally aging counterpart, senescence resistant (SAMR1). While we opted for a male cohort for the P301L dataset to minimize confounding effects of the female oestrous cycle, we here decided to consider female SAMP8 mice given the document stronger impact of aging on microglia in females compared to males (Mouton et al., 2002).

In contrast with some earlier studies (Liu et al., 2020), we could not validate the presence of phospho-tau (AT8) or amyloid pathology via immunohistochemistry or western blot in mice up to 9 months old (Supplementary Fig. 6A-E). Nonetheless, we could confirm compromised hippocampus-dependent spatial learning and memory in the Morris Water Maze (MWM), diminished exploration in the Y-maze, and reduced anxiety, compared to SAMR1 mice evidenced by increased time spent in the open areas of the elevated plus maze and open field tests (Supplementary Fig. 7A-F). While SAMP8 mice demonstrate more pronounced deficits compared to SAMR1 mice, the latter did not show increased activity in the “novel” arm during the Y-Maze probe, suggesting a potential exploratory deficit which may be linked to their advanced age (Supplementary Fig. 7D). In line with the behavioural defects, SAMP8 mice displayed signs of neurodegeneration in the hippocampus and dendate gyrus, such as fewer NeuN-positive neurons and more TUNEL-positive nuclei, which are indicative of apoptosis (Supplementary Fig. 6F-I).

We characterized the microglial heterogeneity at 2, 5 and 9 months for the SAMP8 and SAMR1 model representing a total population of 47,200 non-proliferating microglial cells after exclusion of non-microglial cells and proliferating microglia (Fig. 5A; Supplementary Fig. 8). We will further refer to this dataset as the SAMP8 dataset. DGE analysis on the pooled microglia of all ages revealed 456 genes with significant p-values, 11 of which showing more than 2-fold change between SAMP8 and SAMR1 mice. Compared to SAMR1 mice, SAMP8 mice showed an upregulation of genes involved in metabolism (*Apoe*, *Ldhb, Slc2a5)* and downregulation of immune response genes (*H2-Ob*, *H2-K2*) (Fig. 5B). Many of these differentially expressed genes were present at all considered ages (Supplementary Fig. 9), suggesting that they do not accompany accelerated aging, but rather that they represent global differences in microglial markup that could still prime for this process.

**Figure 5.**
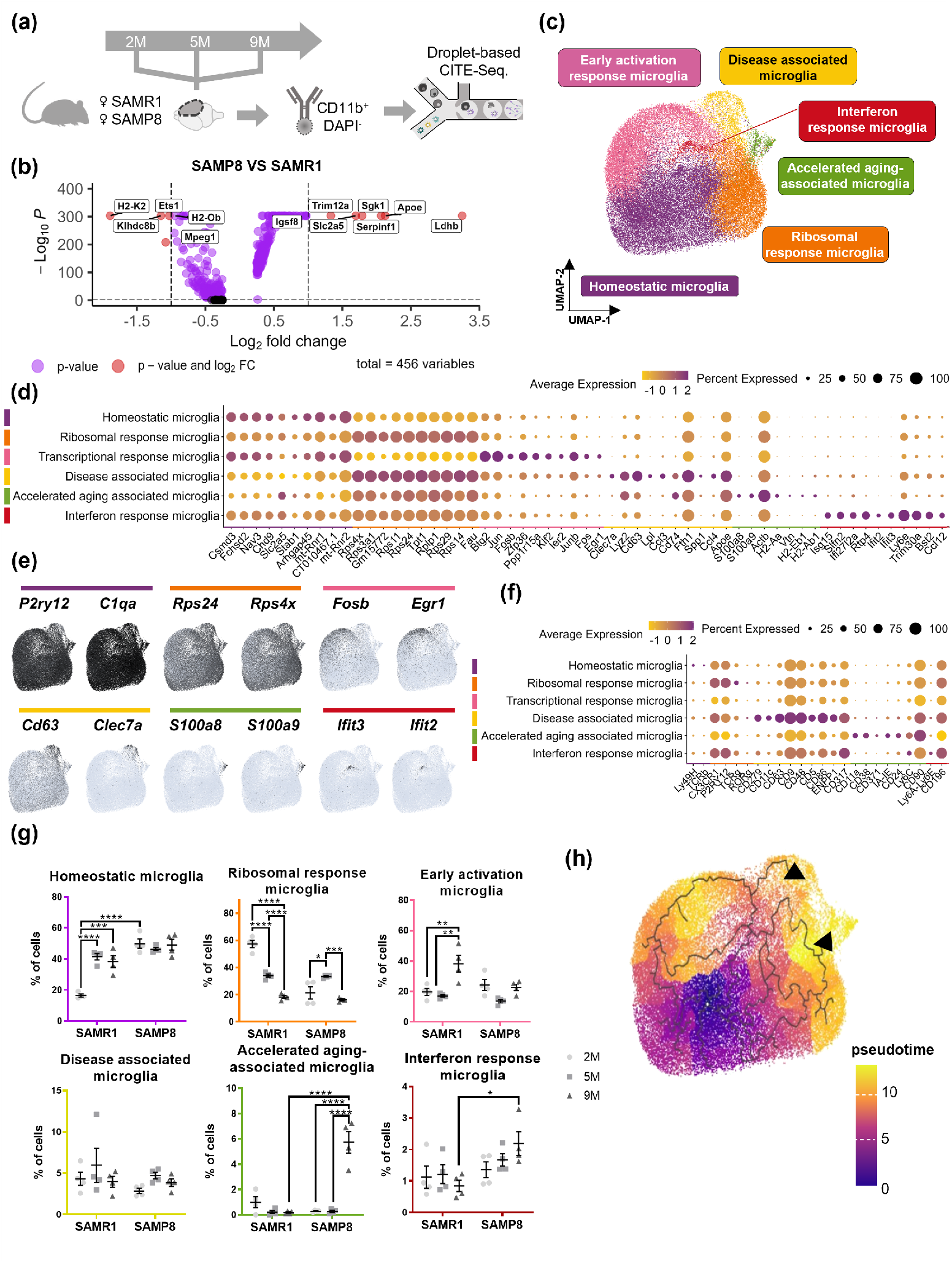
Aging microglia characterized by the high expression of *S100a8* and *S100a9* are specifically present in SAMP8 mice at 9M. (**a**) Schematic diagram showing the CITE-Seq timeline. (**b**) Volcano plot illustrating the genes differentially expressed in microglia between SAMP8 and SAMR1 mice. (**c**) UMAP plot depicting the different microglial subpopulations among all different groups (47200 cells). (**d**) Dot plot representing the top 10 differentially expressed genes of each microglial subpopulation. (**e**) Feature plot depicting the relative expression of two representative genes of each microglial cluster. (**f**) Dot plot displaying the representative protein markers of each microglial subpopulation determined by CITE-Seq. (**g**) Quantification of the percentage of cells in each microglial subpopulation in the different mice and at different time points. (*n* = 4 mice/condition) (**h**) Pseudotime trajectory demonstrating the shift in microglial state. The black arrowheads indicate two separate tracks that end in the aging-associated microglia population. Values are mean ± SEM. Statistical differences (*P < 0.05, **P < 0.01, ***P < 0.001 and ****P < 0.0001) were determined by two-way ANOVA with Tukey’s multiple comparisons test.

### Identification of an Accelerated Aging-Associated Microglial Subpopulation in SAMP8 Mice

Having observed limited change in global microglial gene expression with age (Supplementary Fig. 9), we next asked whether there might be more subtle alterations in microglial diversity in SAMP8 mice. After dimensionality reduction and UMAP representation, we identified the same five microglial subpopulations as also retrieved in the P301L dataset (Fig. 5C-F). Surprisingly, SAMR1 mice exhibited relatively large shifts in different microglial subpopulations with age. At 2 months, they displayed a predominant ribosomal phenotype, but at older ages, a shift towards homeostatic and early activation states was observed. In contrast, SAMP8 mice retained a more stable distribution with only a transient surge in the ribosomal subpopulation at 5 months. However, at 9 months, SAMP8 mice developed a distinct microglial subpopulation that was not seen in SAMR1 mice (Fig. 5G). This previously unidentified subpopulation had some correspondence in gene expression with the DAM phenotype (*e.g.,* upregulation of *Apoe*, *Lyz2* and *Cd74*). Pseudo-time analysis confirmed the closer molecular relationship with DAM (as opposed to homeostatic microglia), but also revealed a connection with ribosomal response microglia (Fig. 5H). When further scrutinizing their unique gene expression signature, we noticed a high expression of major histocompatibility complex class II (MHC-II) related genes (*H2-Aa*, *H2-Eb1* and *H2-fAb1)* and the calprotectin-encoding genes *S100a8* and *S100a9*. Given their specific occurrence in older SAMP8 mice and their characteristic pro-inflammatory marker panel that is related with aging, we coined these microglia *accelerated aging-associated microglia* (A3M) (Frank et al., 2006; Swindell et al., 2013).

### A3M are enriched in K18-P301L mice

The A3M subpopulation exhibited high expression of *S100a8* and *S100a9*, and overlapping gene expression patterns with DAM. We hypothesized that this subpopulation may also be present in the K18-P301L mice, but that it might have been obscured in the analysis due to the concurrent rise in DAM and the overall surge in *S100a8* and *S100a9* expression in this model. Due to the sex differences between the SAMP8 and P301L datasets, complete integration was not possible. Consequently, we defined an A3M-specific gene signature of 30 genes through DGE analysis between the A3M and the DAM subpopulation of the SAMP8 mice scored its enrichment in the P301L dataset. We observed a marked enrichment uniquely in the K18-P301L group relative to the PBS-P301L group (P-value = 0.0181) at 3M (Fig. 6A), suggesting that this population was indeed present and hidden in the dataset. To confirm the presence of these cells *in situ*, we performed immunostaining of contralateral hemisphere sections using S100A8 as distinctive biomarker. Both PBS- and K18-injected P301L mice showed S100A8 predominantly in astrocytes and neurons, but also a distinct microglial subset marked by co-localization of S100A8 and IBA1 (Fig. 6C). This specific microglial population, characterized by a ramified (rather than an amoeboid) morphology (Fig. 6B) was consistently found in the isocortex (in particular in the entorhinal cortex) of the K18-P301L model and it was not observed in the PBS-P301L group (P = 0.0006) (Fig. 6D). In addition, we observed an unusually strong S100A8 signal in a subregion of the hippocampus of P301L+K18 mice, but this signal was not restricted to microglia (Fig. 6C). We confirmed the microglial nature of the S100A8^+^IBA1^+^ double-positive cells through additional co-localization with P2RY12 (Fig. 6B). Despite the observed increase in *S100a9* transcripts, we could not confirm upregulation of S100A9 anywhere in the brain (Fig. 6E), This discrepancy might be attributed to the antibody’s inability to recognize the epitope, potentially due to differing conformations or oligomer formation (S. Wang et al., 2018).

**Figure 6.**
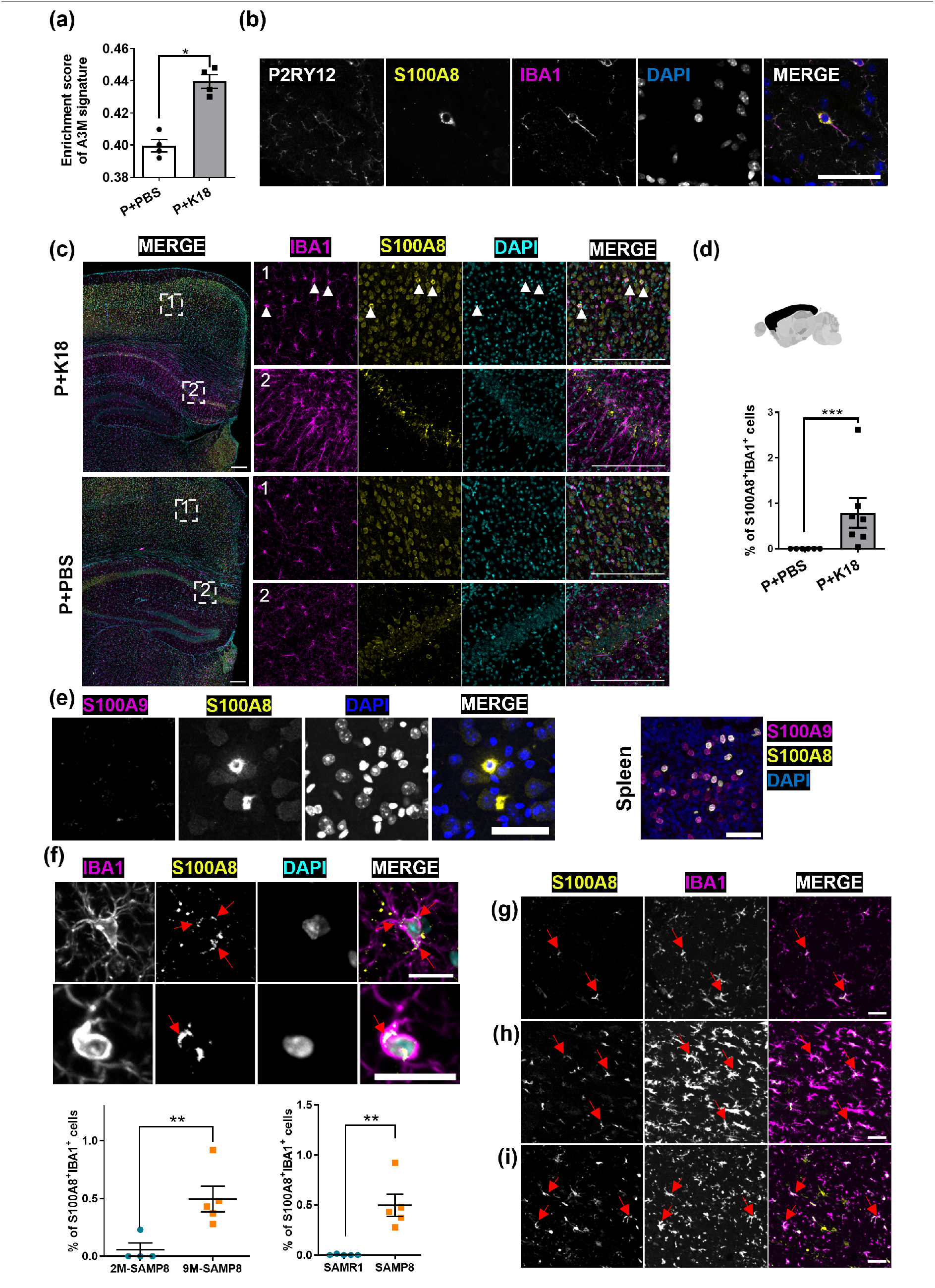
S100A8^+^IBA1^+^ cells are present in mice and human samples showing aging, AD or tauopathy. (**a**) Average enrichment score of the A3M signature in FVB and P301L-injected mice at 3 months post-injection, derived from CITE-seq data. The A3M signature is defined by the top 30 genes differentiating A3M from disease-associated microglia (DAM) in SAMP8 mice. Enrichment in individual cells was determined using the AddModuleScore function, which evaluates the collective expression of this specific gene set. (**b**) Representative images of P2RY12 staining of the S100A8^+^IBA1^+^ positive cells. (**c**) Representative image of IBA1, S100A8, and DAPI staining in P+PBS and P+K18 mice. Montages on the right show a zoom of the delineated region presented on the left image. The first square (1) represents a zoom in the cortex where S100A8^+^IBA1^+^ cells (white arrowheads) were localized in P+K18 mice. The second square (2) represents a zoom in the hippocampus where an intense S100A8 signal was observed in the P+K18 mice compared to P+PBS mice. Scale bar 200 µm. (**d**) Quantification of the percentage of S100A8^+^IBA1^+^ cells over the total number of IBA1^+^ cells in the isocortex at 3-months post-injection in PBS-(P+PBS) and K18-(P+K18) injected P301L mice (1 slice/mouse). (**e**) Representative images showing S100A9 staining of S100A8^+^IBA1^+^ cells in the brain (left part) and S100A8, S100A9, DAPI immunostaining in the spleen (right part), using the same antibodies, serving as a positive control. Scale bar 50 µm. (**f**) Representative, contrast-enhanced images of two S100A8+IBA1+ cells (upper section) and quantification (lower section) of S100A8^+^IBA1^+^ positive cells over the total number of IBA1^+^ cells in the hippocampus of SAMP8 and SAMR1 mice at the age of 9 months (1 slice/mouse, *n* = 5-6 mice/condition). Red arrows show intracellular staining. Scale bar = 20 µm. (**g-i**) Representative image of S100A8^+^IBA1^+^ double positive staining (red arrows) of an aged non-neuropathologic patient (69 years old) (**g**), AD (**h**) and Tauopathy (**i**) diagnosed patients. Scale bar 50 µm. Values are mean ± SEM. Statistical differences (\**P* < 0.05, \*\**P* < 0.01, \*\*\**P* < 0.001) were determined by non-parametric one-tailed Mann-Whitney U test.

S100A8^+^IBA1^+^ microglia were also identified in brain sections of SAMP8 mice at 9 months old but not in their younger, 2-month-old counterparts, nor in SAMR1 mice at 9 months (Fig. 6F). Surprisingly, the intensity of S100A8 staining was weaker in SAMP8 mice compared to K18-P301L mice, and the S100A8-positive cells were predominantly found in the hippocampal region (more specifically in the stratum oriens, statum radium and stratum lacunosum-moleculare). Finally, the lack of additional internalized nuclei suggests that the signal is not likely associated with phagocytosis. Thus, both 3-months p.i. K18-P301L mice and 9-month-old SAMP8 mice exhibit a distinct microglial phenotype, characterized by elevated levels of S100A8 but with different penetrance and preferential location.

### S100A8-enriched microglia are also present in human brain

To assess the presence of this microglial subpopulation in humans, we analyzed post-mortem brain tissues from both male and female patients, aged 51 to 70, diagnosed with tauopathies or AD, as well as from control individuals aged 41-69 with no neuropathological diagnoses. We concentrated on brain sections encompassing the entorhinal region and hippocampus, correlating with the observed locations of these cells in K18-P301L and SAMP8 mice, respectively. S100A8^+^ microglia were identified in 3 out of 10 AD patients aged 57, 53, and 62 years, in 1 out of 10 patients with tauopathies aged 51 years, and in 1 control individual aged 69 years, situated within the hippocampus and/or regions encompassing the parasubicular, subicular, and entorhinal cortices (Fig. 6G-I). This evidence suggests that the identified subpopulation is not exclusive to murine models and is not solely linked to tau pathology. The detection of these cells in only one elderly individual without neuropathological diagnosis points to their potential connection with accelerated aging, but more comprehensive studies are warranted.

## Discussion

In this work, we have illustrated the diversity and plasticity of microglial phenotypes and a conspicuous emergence of S100A8^+^ microglia in both accelerated aging and tauopathy. First, we confirmed that depletion of microglia can dampen the hyperphosphorylated tau load in the brain. This supports the unintentional adverse role of microglia in hyperphosphorylated tau spreading, as previously suggested by other studies (Asai et al., 2015; Chen et al., 2023; Mancuso et al., 2019; C. Wang et al., 2022). Hyperphosphorylated tau accrual was blunted in specific brain regions, but the correlation with changes in microglial density or morphology was incomplete. Even within the cortex, a region with a homogenous microglial occupation and morphology, we found compartments that were more affected by microglial depletion than others. This region-dependency made us wonder whether specific microglial subpopulations drive tau pathology progression.

CITE-Seq did not reveal a subpopulation unique to the K18+P301L model, but we did observe a modest increase in the DAM population, consistent with other single-cell RNA sequencing studies in other tauopathy models, such as TE4 mice (P301S expressing human APOE4) and Tau4RΔK-AP mice (Chen et al., 2023; Kim et al., 2022). In line with the Tau4RΔK-AP mice study, the increase of the DAM subpopulation was only observed at late stages of hyperphosphorylated tau spreading (Kim et al., 2022). Furthermore, there was no one-on-one relationship between DAM emergence and hyperphosphorylated tau load. Using CD63, we localized the DAM increase in P301L+K18 brain slices to the dorsal region of the brain, which includes the brainstem, corticospinal tract and the striatum. We found a clear impact of microglia depletion on hyperphosphorylated tau load in the striatal dorsal region, but not in other DAM-harbouring regions such as the thalamus or brain stem. Conversely, hyperphosphorylated tau load in the isocortex was sensitive to PLX treatment but did not harbour any putative DAM. Thus, while it is tempting to speculate that the selective persistence of DAM may be responsible for the enhanced spreading, their low numbers and lack of correlation with sites of hyperphosphorylated tau, suggest that DAM are not the main driving factor behind the spreading process, but rather a bystander or target thereof. Given that PLX3397 can also affect the peripheral immune system and that monocyte-derived macrophages with an M2-like profile have been implicated in tauopathy mouse models as well (Ben-Yehuda et al., Molecular Neurodegeneration, 2021), one possibility is that the observed regional tau reduction may be attributed to the concomitant depletion of the macrophage pool. This could account for the lack of observed correlation between tau spreading and the evolution of microglial phenotypes in our study. Another possibility is that the activity and response of common microglial subpopulations to tau adds to the spreading process without directly affecting their function.

Redirecting our attention to the population-level gene expression changes, we noticed in 6-months-old P301L+K18 mice (3 months p.i.), an upregulation of the calprotectin-encoding genes, *S100a8* (also termed *MRP8* or calgranulin A) and *S100a9* (also termed *MRP14* or calgranulin B). Interestingly, studies have found that the co-expression of *S100a8* and *S100a9* represents a robust feature of brain aging, as both aged human and mouse brains show higher expression levels compared to younger samples (Swindell et al., 2013). The genes have been inferred in AD development as well. Calprotectin levels are elevated in the brain of AD patients and mouse models for amyloidosis (Kummer et al., 2012; Lodeiro et al., 2016), and its depletion reduces amyloid burden (Ha et al., 2010; Kummer et al., 2012). In PS/APP mice, the S100A8/A9 heterodimer is upregulated in microglial cells surrounding amyloid plaques (Kummer et al., 2012). However, the association with tau pathology is less clear. S100A9 is present in activated glia and neurons positive for neurofibrillary tangles (Shepherd et al., 2006), but an inverse correlation between S100A9 and hyperphosphorylated tau immunostaining was reported in neurons (C. Wang et al., 2014). This underscores the weak causal relationship and places calprotectin downstream of tau pathology.

Supporting this, we discovered that the upregulation of exactly this gene set typified a unique microglial population in aged SAMP8 mice. While some studies have reported AD hallmarks such as amyloid or tau pathology in this model (Liu et al., 2020), we could not replicate this using immunofluorescence staining or western blot suggesting that their levels (if present) are below those of most advanced AD models and thus do not represent a major contributor to or proxy of the aging process. Nevertheless, consistent with other studies on SAMP8 (Liu et al., 2020), we noted higher spatial learning and memory deficits, reduced anxiety, and signs of neurodegeneration in SAMP8 compared to SAMR1 mice (although the latter showed lower performance on the Y-Maze test than expected).

The A3M subpopulation that we discovered in SAMP8 mice shares some similarities with DAM, including high expression of *Cd74*, *Apoe* and *Lyz2.* However, this subpopulation expresses a specific gene pattern that sets it apart from other microglial subpopulations. Along with the high level of calprotectin protein, we also found increased expression of MHC-II related genes such as *H2-Aa* and *H2-Eb1* in this subpopulation. MHC-II, which is responsible for presenting processed antigens to immune cells, has been shown to increase during aging in microglia (Norden & Godbout, 2013; Sheffield & Berman, 1998). Interestingly, this subpopulation also shares some similarities with the Late-stage AD-Associated Microglia (LADAM) described in the Tau4RΔK mice (Kim et al., 2022). LADAM subpopulations express high levels of MHC and S100 family genes as *Cd74*, *H2-Eb1*, *H2-Ab1*, *H2-A2*, *S100a4*, *S100a6*, and *S100a10*. Additionally, the LADAM subpopulation was described to be derived from Early-stage AD-Associated Microglia (EADAM), which show high levels of interferon relative genes *Irf7* and *Isg15*, or from a DAM2 subpopulation characterized by high levels of *Spp1*, *Gpnmb*, *Apoe*, *Cst7*, *Cd74*, *Lpl*, and *Lgals3* (Kim et al., 2022). Interestingly, except for *Gpnmb*, all DAM2-related genes were expressed by the DAM subpopulation in our data set, suggesting that the A3M subpopulation is a possible descendent from the DAM lineage, rather than the ribosomal subpopulation.

An important limitation of our study is the difference in gender balance between both models (P301L and SAMP8). Although this hindered a direct comparison and integration of both datasets, we could still detect the enrichment of A3M in K18-P301L mice by projecting their characteristic gene set. This was further confirmed by immunostaining. Thus, although we cannot yet conclusively establish that the A3M identified in both models represent the exact same subtype, our initial findings suggest they are alike. The detection of the S100A8^+^IBA1^+^ microglia subpopulation in human brain tissue further confirms that this subpopulation also exists in the human brain. Interestingly, within the samples that were positive, we found younger AD (53y) or Tau (51y) patients than the positive healthy patient (69y), which may indeed corroborate their putative correlation with accelerated brain aging.

Overall, our findings suggest that the upregulation of S100A8 in microglia is a common feature of accelerated aging and tau pathology, and that tau pathology can evoke their emergence at an earlier age, suggesting they accelerate the brain aging process. Future research should investigate the role of A3M in AD development and the link between tauopathy and accelerated aging.

## Acknowledgements

We are thankful to the VIB Flow Core Ghent, VIB Single Cell Core and VIB Nucleomics for support and access to the instrument park (vib.be/core-facilities). Additionally, we are grateful to the NeuroBiobank of the Born-Bunge Institute (IBB-Neurobiobank), Wilrijk (Antwerp), Belgium (ID:BB190113) for their assistance and provision of human tissue samples. We would also like to extend our gratitude to Elien Theuns and Karen Sterck for their invaluable assistance in conducting additional experiments.

## Conflict of Interest statement

The authors report no competing interests.

## Funding statement

This work is supported by the Alzheimer Foundation SAO-FRA (Grant Nr. 2019/0035), VLAIO [SynDAM, Grant Nr. HBC.2019.2926], Research Foundation Flanders [FWO Grant Nr. I003420N and IRI I000123N] and the University of Antwerp [BOF IMARK, µNEURO, IOF FFI210242].

## Author’s contributions

The authors confirm contribution to the paper as follows: study conception and design: W.H.D.V., R.G.; conducted the experiments and acquired the data: R.G., J.V.D.D., S.T., R.W., D.W., D.V.D., R.V., N.V.; analysis and interpretation of results: R.G., B.B., R.M., J.D.P.A., P.V., J.R., D.V.D.; draft manuscript preparation: W.H.D.V., R.G., J.V.D.D., P.V., R.V., N.V., B.B., R.M., J.R., D.V.D. All authors contributed to the article and approved the submitted version.

## Data availability

Our image analysis scripts are available on Github (github/devoslab).

We are in the process of submitting our CITEseq datasets to NCBI and will communicate the link once they have become available.

## Notes

### Competing Interest Statement

The authors have declared no competing interest.

