## Supplementary data for "S100A8-enriched microglia populate the brain of tau-seeded and accelerated aging mice"

### Supplementary Materials and methods

#### Tunel staining

For the detection of apoptosis in situ, tissue sections were subjected to the Click-iT™ Plus TUNEL Assay for In Situ Apoptosis Detection (C10619, Invitrogen), utilizing the Alexa Fluor™ 647 dye according to the manufacturer's protocol. Fresh PFA-fixed brains were dissected into coronal 30 µm sections. These sections were then re-fixed with paraformaldehyde, permeabilized with proteinase K, and then incubated with TdT enzyme for the labeling of free 3'-OH termini in DNA fragments, a hallmark of apoptosis. Following the labeling step, the sections were exposed to the Click-iT™ Plus reaction cocktail containing the Alexa Fluor™ 647 azide, which specifically reacts with the TdT-mediated dUTP incorporated in the nicked DNA ends via a copper-catalyzed click reaction. After thorough washing to remove unbound components and staining with DAPI (5 µg/ml), the stained tissue sections were mounted with a coverslip using Citifluor™ Mounting solution AF-1. Confocal images were acquired of the slices (as tile scan) with a Nikon Ti2 W1 spinning disk confocal using a 20x/NA 0.75, stitched, flattened and quantified with QuPath as described in previously in the material & method section of the article.

#### Immunochemistry staining on slices

After perfusion and PFA 4% fixation, brains were embedded in paraffin and sliced into sagittal 5 µm sections using microtome (Leica). Following antigen retrieval with 70% formic acid for 10 minutes at RT and endogenous peroxidase and biotin/avidin blocking, the tissue was incubated with AG8 antibody (1:2000, Biolegend, 800704) overnight at RT. For detection, the HRP conjugated streptavidin from Invitrogen and DAB from Dako (K3468) were used and counterstained with hematoxylin. For the Alcian blue-Periodic Acid Schiff (PAS), we used the kit from Abcam (ab245876), according to the manufacturer recommendation. Images were acquired on Axiophot (Zeiss, 20x/NA 0.60, pixel size: 0.16 x 0.16 $\mu m^2$).

#### Total RNA and protein extraction, retrotranscription and quantitative real-time RT-PCR

Total RNA was extracted from CD11b-positive cells using Qiagen RNeasy mini kit (74104, Qiagen). RNA was quantified using NanoDrop 2000 spectrophotometer (Thermo Fisher Scientific) and retrotranscription (250 ng of total RNA) was done using iScript™ cDNA Synthesis Kit (1708891, BioRad) and T100 Thermal Cycler (BioRad). RT-PCR was performed using SsoAdvanced Universal SYBR® Green (1725272, BioRad) using an CFX Connect Real-Time PCR System (BioRad) with primers for the following genes of interest: *Dusp1*, *Egr1*, *Fos*, *Fosb*, *Nfr4a1*, *Tnfa*, *Il1b* (Supplementary Table 5).

#### Western blot

Snap-frozen brains, removed from olfactive bulbs and cerebellum, were thawed on ice in Buffer H (pH 7.6) containing 10 mM Tris-HCL, 0.8 M NaCl, 10% sucrose, 1mM EGTA containing 1X Halt™ Protease and Phosphatase Inhibitor Cocktail (Thermo Fisher Scientific, 78440) and lysed using Precellys lysing kit (P000918-LYSK0-A, Bertin Technologies) and Minilys homogenizer (Bertin Technologies). Surpernatant was collected after centrifugation at 16,400 g during 40 min at 4°C for western blot. Protein concentration was measured with the Pierce™ BCA Protein Assay Kit (Thermo Fisher Scientific, 23227). Tissue lysates were mixed with 25% NuPage LDS sample buffer (Thermo Fisher Scientific, NP0007) and 5% dithiothreitol (DTT, Thermo Fisher Scientific NP0009) and heated for 5 minutes at 95°C. 80 µg of each sample was loaded onto NuPAGE™ Novex 4–12% Bis-Tris Protein Gels (Thermo Fisher Scientific, NP0322PK2), with MOPS SDS running buffer (Thermo Fisher Scientific, J00047). PageRuler™ Prestained Protein Ladder was used as marker (Thermo Fisher Scientific, PI26617). Next, proteins were transferred to nitrocellulose membrane (Invitrogen, LC2000) at 30V during 1h, using a transfer mixture of NuPAGE transfer buffer (Thermo Fisher Scientific), NuPAGE antioxidant (Thermo Fisher Scientific) and methanol. Afterwards the membranes were blocked in blocking buffer (5% ECL (Sigma GERPN418) in Tris Buffered Saline with 0.2% Tween 20 (TBST)), and subsequently incubated with primary antibodies AT8 (1:2000, mouse pSer202/Thr205/PSer208, produced at Janssen Pharmaceutica), diluted in blocking buffer. Anti-beta actin antibody (1:2000, mouse, ab8226, Abcam) was used as a reference protein. Horseradish peroxidase (HRP)-conjugated goat anti-mouse (Sigma-Aldrich A4416, 1/2000) was used as secondary antibodies. Proteins were detected by chemiluminescence with ClarityTM western ECL substrate (Bio-Rad, 170-5060) using a western blot Imager (Bio-Rad, ChemiDocTM Touch Imaging System) and analyze using ImageJ (Schneider, Rasband, & Eliceiri, 2012).

### Behavioural tests

Behavioural deficits of SAMR1 and SAMP8 mice at the age of 9 months were assessed using Morris Water Maze (MWM), Y-Maze, elevated plus maze and open-field test at the Laboratory of Neurochemistry & Behaviour (Wilrijk, Belgium).

#### Morris Water Maze

The MWM assesses hippocampus-dependent visuo-spatial learning and memory (Hendrickx et al., 2022). The setup consisted of a circular pool (diameter: 150 cm, height: 30 cm) filled with opacified water using non-toxic white paint and was kept at 25 °C. Invariable visual cues were placed around the pool. The MWM consisted of an acquisition phase and a probe trial. The acquisition phase was performed over a period of 4 days and consisted of 2 daily trial blocks (at 10:30 AM and at 03:00 PM) of 4 trials with a 15 min inter-trial interval. During the acquisition phase, a round acrylic glass platform (diameter 15 cm) was placed 1 cm below the water surface in a fixed position in the centre of one of the pool's quadrants. Mice were placed in the water facing the wall and were recorded while trying to find the hidden platform for a maximum duration of 120 s. If the mouse was not able to reach the platform within 120 s, it was guided to the platform, where it had to stay for 15 s before being returned to their home cage. The starting positions varied in a semi-random order. The probe trial followed 4 days after the final acquisition trial. For this trial, the platform was removed, mice were placed in the MWM at a fixed position, and swimming trajectories were recorded for a period of 100 s. During both acquisition and probe trials, the animals' trajectories were recorded using a computerized video-tracking system (Ethovision), with path length, escape latency, and swimming speed recorded.

#### Y-Maze

Y-Maze Spontaneous Alternation is a behavioural test to measure the willingness of rodents to explore new environments. Rodents typically prefer to investigate a new arm of the maze rather than returning to one that was previously visited. It is used to investigate exploratory behaviour and cognitive function related to spatial working memory. Testing occurred in a Yshaped maze with three white, opaque plastic arms (length: 33 cm, width: 5 cm) at a 120° angle from each other. After introduction to the centre of the maze, the animal is allowed to freely explore the three arms. Over the course of multiple arm entries, the subject should show a tendency to enter a less recently visited arm. On day 1, the mouse was allowed to freely explore the maze for 8 min, while on day 2, one arm was closed off and the animal was allowed to explore for 10 min (trial run). After 4h, the animal was subjected to a probe trial where it was allowed again to explore the complete maze (5 min). The number of arm entries and the number of triads were recorded (Ethovision, Noldus, Wageningen, The Netherlands) to calculate the percentage of alternation. An entry occured when all four limbs were within the arm.

#### Elevated plus-maze

The elevated plus-maze apparatus was used to evaluate anxiety (Vloeberghs, Van Dam, Franck, Staufenbiel, & De Deyn, 2007). It consisted of four cross-shaped arms, of which two were brightly lit and open, and the other two dark and enclosed (30 cm × 5 cm × 15 cm; length × width × height of the enclosed arms). The maze itself was elevated 60 cm from the floor and mice were always placed in the central area (5 cm × 5 cm), facing the left enclosed arm. All mouse movements were tracked by camera for 5 min (Ethovision). The number of entries, latencies to the first arm entry, the duration of the time spent in open and closed arms, the total distance moved, and velocity were the parameters measured.

#### Open-field

Open-field behaviour was measured for 10 min during the dark phase of the animal's activity cycle in a brightly lit 50×50 cm2 arena (Van Dam et al., 2003). Mice always started from the same corner of the arena and were allowed 1 min of acclimation to the setup before recording. A computerized video tracking system (Ethovision) was used to record trajectories and calculate path length and number of entries in the centre circle or the 7 × 7 $cm^2$ corners of the open field.

### Supplementary Figures

**Supplementary Figure 1. Actinomycin D and Brefeldin A treatment reduce microglia activation during neural dissociation.** The fold change of early immediate genes (**a**) and pro-inflammatory cytokine genes (**b**) expression was measured using RT-PCR after brain dissociation and Percoll gradient in the absence (CTRL) or presence of Brefeldin A (1x) and Actinomycin D (5 nM) (+BrefA +ActD). Values are mean ± SEM. Statistical differences (*$P$ < 0.05) were determined by nonparametric one-tailed Mann-Whitney U test. ($n$ = 4-5 mice/group)

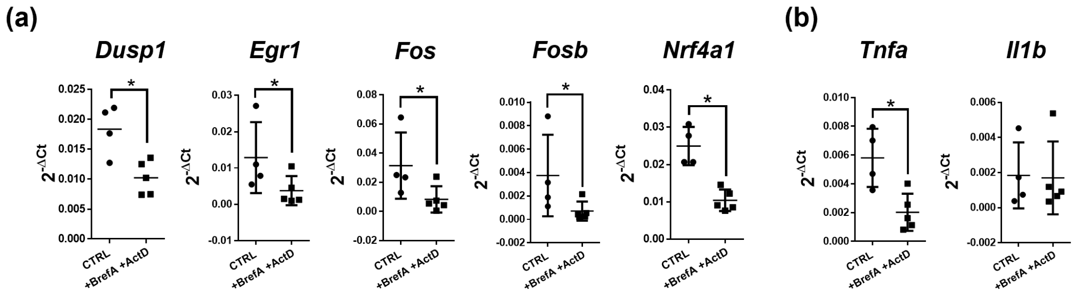

**Supplementary Figure 2. PLX3397 depletes microglia significantly and more homogeneously in CX3CR1$^{+/GFP}$ mice compared to PLX5622.** (**a**) Schematic diagram showing the microglial depletion strategies of CX3CR1$^{+/GFP}$ mice. Flow cytometry plots illustrate the gating strategy for assessing single-cell microglia based on CX3CR1-GFP$^{+}$ signal. (**b**) Quantification of microglial proportion, represented by the percentage of CX3CR1-GFP$^{+}$ cells relative to the total single-cell population. (**c**) Representative images of CX3CR1-GFP hemisphere signal with a zoom. Scale bar 400 µm. (**d**) Quantification of the microglia density in brain slices in the different treated groups. More than 70% of microglia was depleted after PLX3397 or PLX5622 treatment (average of 2 slices/mice). (**e-g**) Quantification of number of neurons, astrocytes and oligodendrocytes using respectively NeuN (**e**), Sox9 (**f**), Sox10 (**g**) staining after 28 days of diet in different brain regions (average of 2 slices/mice). (**h**) Representative images of coat discoloration after PLX3397 treatment after 28 days of treatment. Values are mean ± SEM. Statistical differences (*$P < 0.05$, **$P < 0.01$ and ***$P < 0.001$) were determined using one-way ANOVA with Dunn's multiple comparisons test for each time-point and each region ($n$ = 2-5 mice/group).

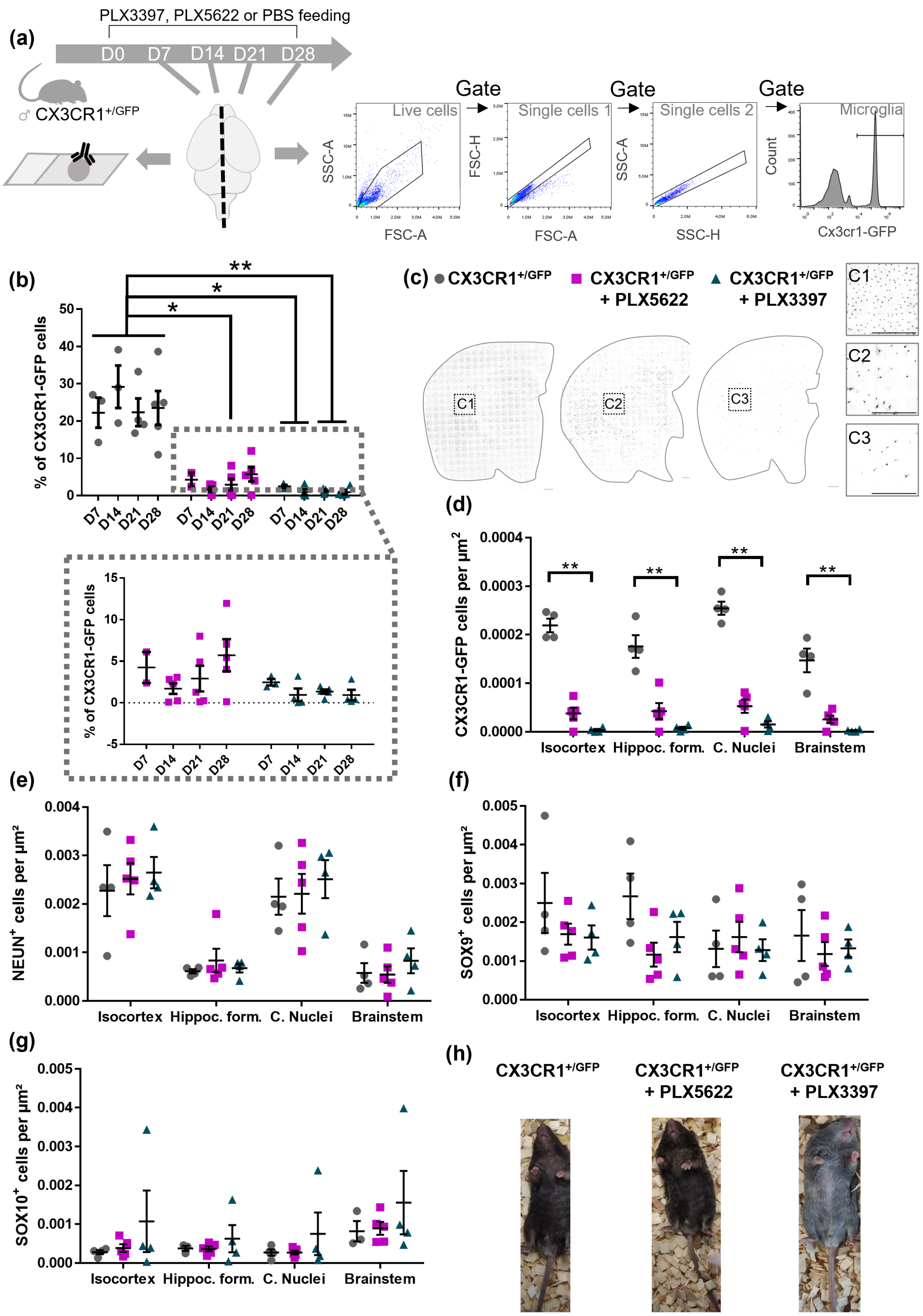
(a)
PLX3397, PLX5622 or PBS feeding
D0 D7 D14 D21 D28
♂ CX3CR1+/GFP
Gate
Gate
Gate
Live cells
Single cells 1
Single cells 2
Microglia
SSC-A
FSC-A
FSC-H
FSC-A
SSC-A
SSC-H
Count
Cx3cr1-GFP
(b)
% of CX3CR1-GFP cells
(c)
CX3CR1+/GFP
CX3CR1+/GFP + PLX5622
CX3CR1+/GFP + PLX3397
C1
C2
C3
(d)
CX3CR1-GFP cells per µm2
Isocortex Hippoc. form. C. Nuclei Brainstem
(e)
NEUN+ cells per µm²
Isocortex Hippoc. form. C. Nuclei Brainstem
(f)
SOX9+ cells per µm²
Isocortex Hippoc. form. C. Nuclei Brainstem
(g)
SOX10+ cells per µm²
Isocortex Hippoc. form. C. Nuclei Brainstem
(h)
CX3CR1+/GFP
CX3CR1+/GFP + PLX5622
CX3CR1+/GFP + PLX3397

**Supplementary Figure 3. PLX3397 efficiently depletes microglia in K18-seeded P301L mice.** (**a**) Schematic diagram showing the microglial depletion strategy. (**b**) Representative images of IBA1 staining in cortex and hippocampus. (**c**) Quantification of microglia was assessed in different regions in PBS-injected FVB (F+PBS), PBS-injected (P+PBS) and K18-injected (P+K18) P301L mice, after PBS (dot) or PLX3397 treatment (square). Values are mean ± SEM (4-5 slices/mouse). Statistical differences (*$P$ < 0.05, **$P$ < 0.01 and ***$P$ < 0.001) were determined by nonparametric one-tailed Mann-Whitney U test for each group. ($n$ = 3 mice/group). Scale bar 100 µm. Cx = Cortex, Hp = Hippocampus, Hippoc. Form = hippocampal formation, C. Nuclei = cerebral nuclei.

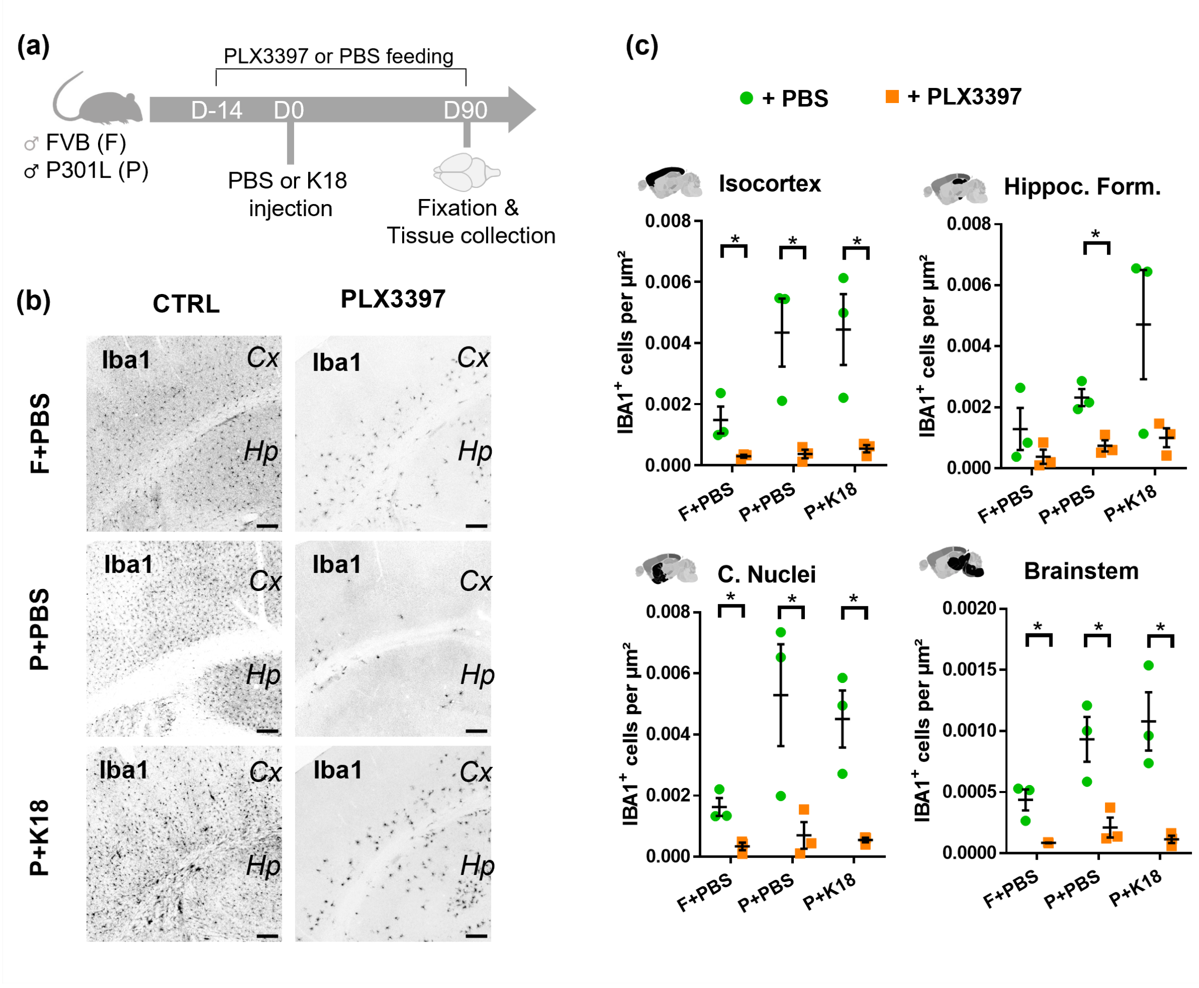

**Supplementary Figure 4. Microglia depletion does not affect tau load in the ipsilateral hemisphere.** (**a**) Representative images of AT8 signal in PBS or PLX3397 treated K18-injected P301L mice. Images represent AT8 staining on a whole sagittal slice and a zoom of a selected region. Scale bar 400µm. (**b**) Quantification of AT8 staining in different regions of the ipsilateral hemisphere after 104 days (14 days pre- + 3 months post-injection) of feeding with modified AIN76A supplemented with PBS (dot) or PLX3397 (square). Positive pixels were quantified in different regions (represented in black) in K18-seeding P301L. Values are mean ± SEM. ($n$ = 7 mice/condition). Hippoc. form. = Hippocampal formation.

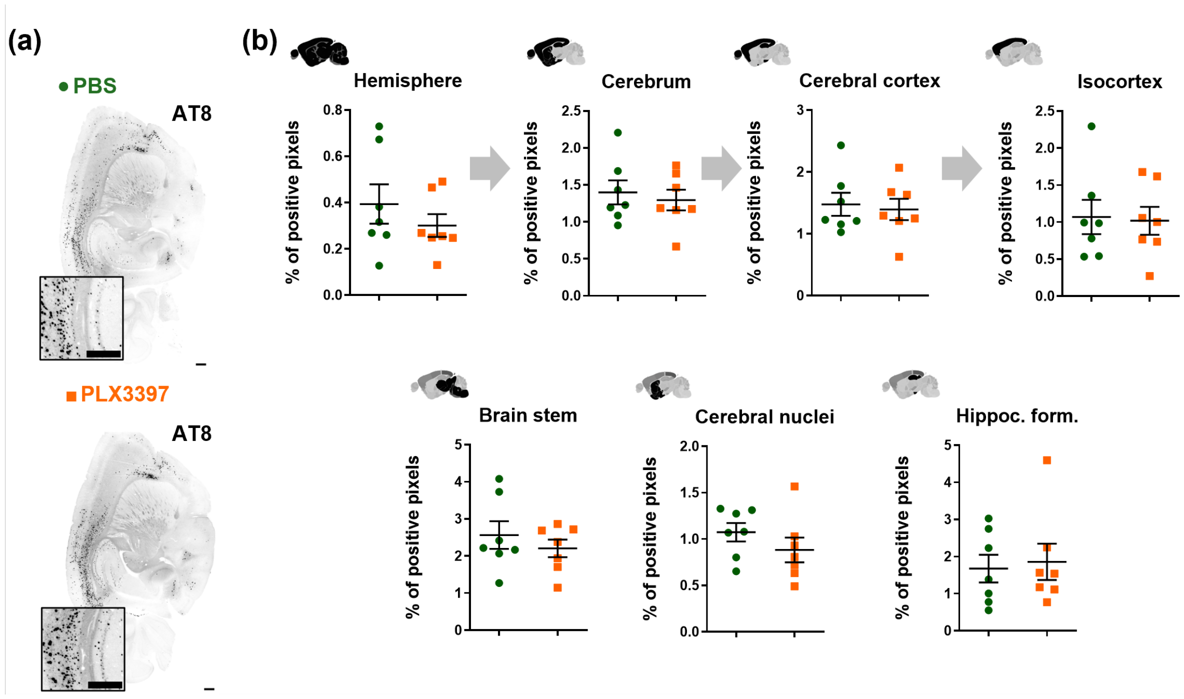

**Supplementary Figure 5. Characterization of isolated CD11b+ cells from FVB and P301L mice.** (**a**) UMAP plot of CD11b+ cells depicting the different cell types of all treated groups (59129 cells). (**b**) Dot plot displaying the representative protein markers of the different cluster. (**c**) Dot plot depicting the top 10 differentially expressed genes of each cluster. (**d**) Feature plot representing specific markers of each cell type.

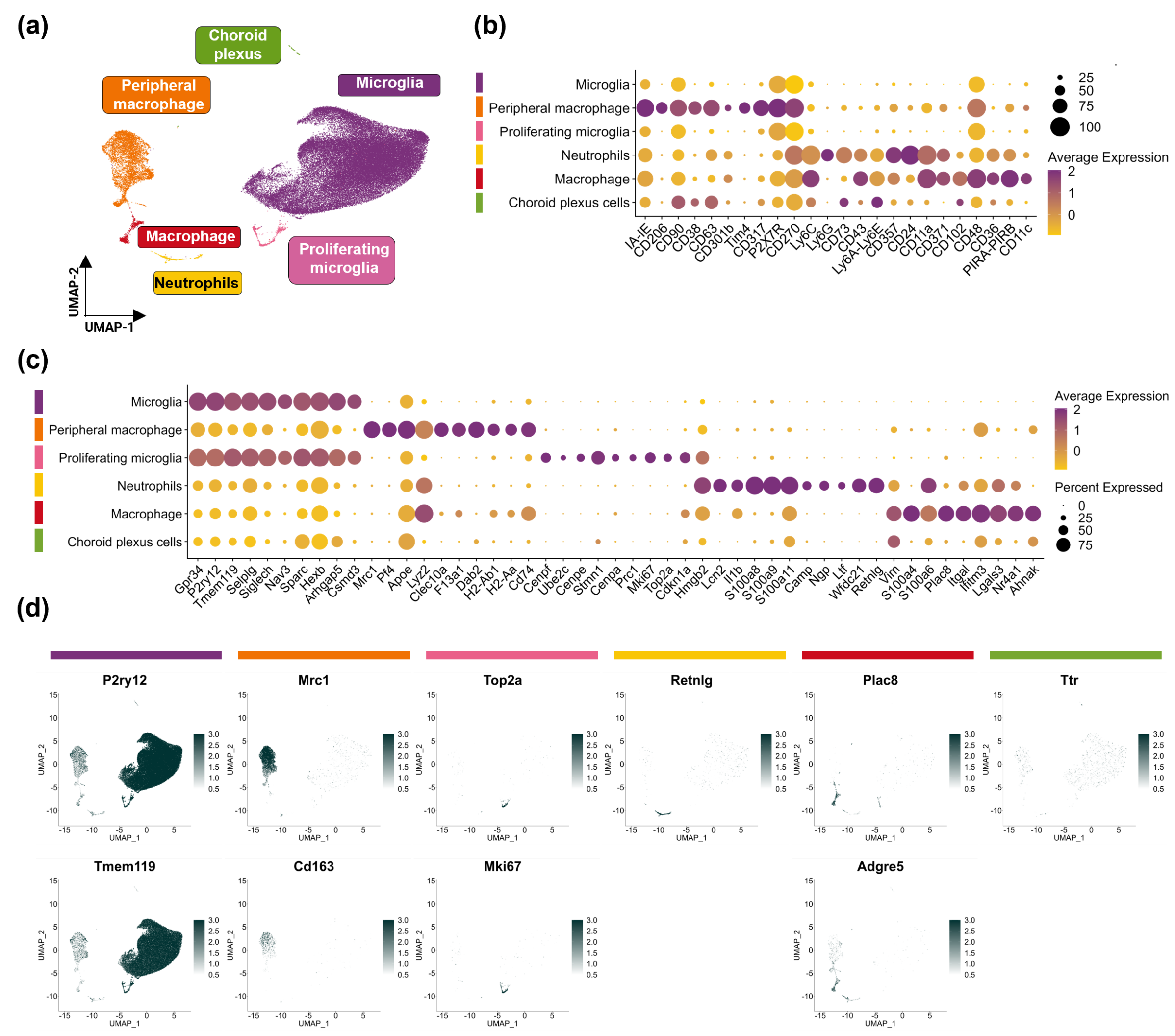

**Supplementary Figure 6. Absence of amyloid and tau aggregates in SAMP8 and SAMR1 brains accompanied by indicators of neurodegeneration.** Representative images of 4G8 (**a**), periodic-acid-schiff (PAS) (**b**), pentameric formyl thiophene acetic acid (pFTAA) (**c**) and AT8 (**d**) staining in brains of SAMP8 and SAMR1 at 9-months old. The last column depicts positive staining of PAS in colon of SAMR1 mice and of AT8 staining in K18-injected P301L mice using similar acquisition parameters. (**e**) Representative western blot images for AT8 and the housekeeping protein beta-actin. Representative immunostaining images (**f**, **i**) with quantification (**g**, **h**) of TUNEL staining per square micrometer of tissue (**f**, **g**) and of the percentage of NeuN-positive cells relative to all DAPI-positive cells in the hippocampus and its subregions (**h**, **i**) (1 image per mouse). Values are mean ± SEM. Scale bar 200 µm.

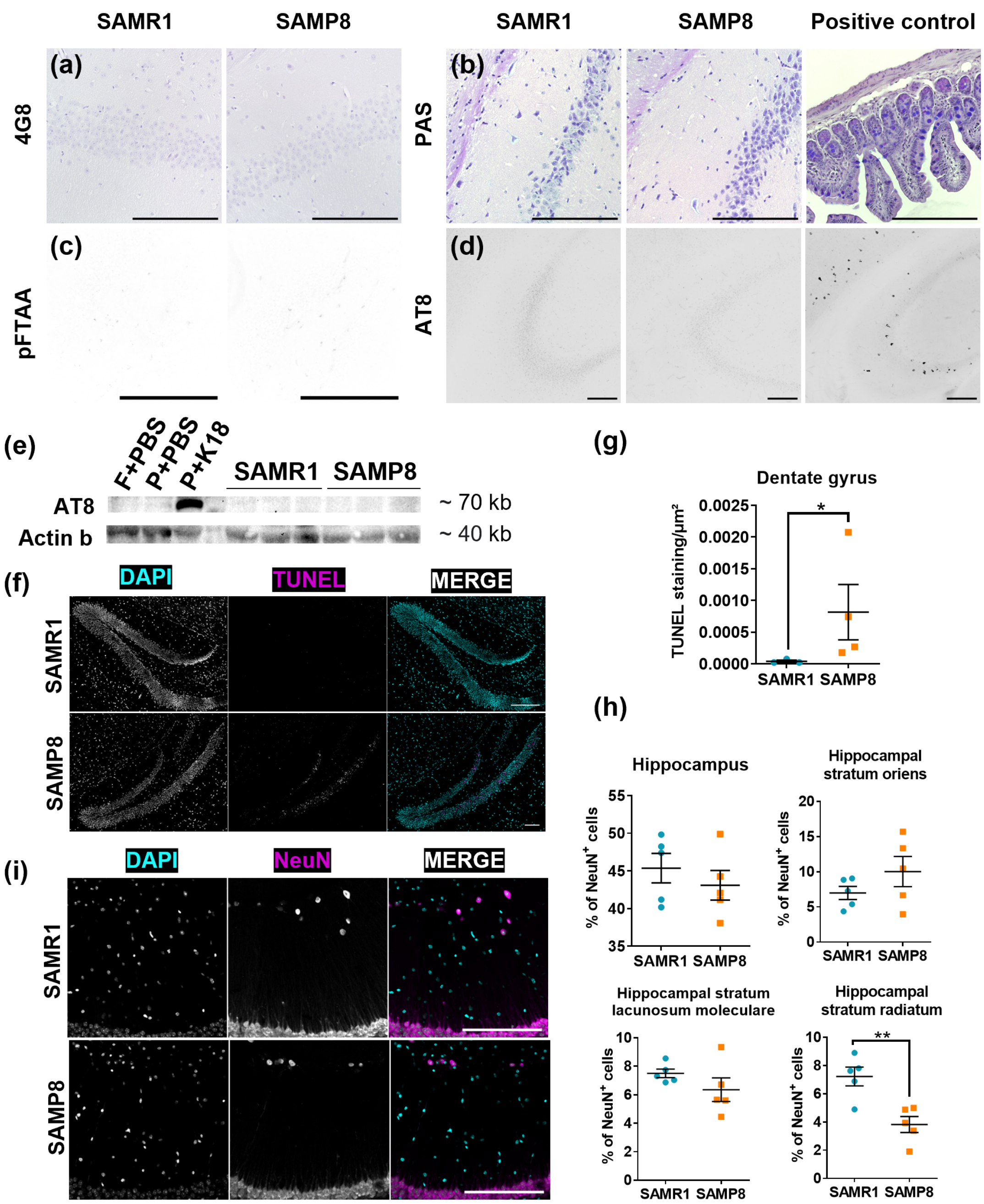

SAMR1
SAMP8
SAMR1
SAMP8
Positive control
(a)
4G8
(b)
PAS
(c)
pFTAA
(d)
AT8
(e)
F+PBS
P+PBS
P+K18
SAMR1
SAMP8
AT8
Actin b
~ 70 kb
~ 40 kb
(f)
DAPI
TUNEL
MERGE
SAMR1
SAMP8
(g)
Dentate gyrus
TUNEL staining/µm²
SAMR1 SAMP8
(h)
Hippocampus
Hippocampal stratum oriens
% of NeuN+ cells
Hippocampal stratum lacunosum moleculare
Hippocampal stratum radiatum
(i)
DAPI
NeuN
MERGE

**Supplementary Figure 7. SAMP8 mice exhibit reduced anxiety and spatial memory and learning impairment at the age of 9 months.** At 9 months, the behavioural results of SAMP8 mice (orange square, n = 10 mice) were compared to those of SAMR1 mice (blue circle, n = 12 mice). Schematic diagram (left) and results (middle and right) of the Morris Water Maze (MWM) trials (**a**), the MWM probe (**b**), the Y-maze trials (**c**), the Y-maze probe (**d**), the elevated plus maze test (**e**), the open-field test (**f**). Values are mean ± SEM. Statistical differences (*$P < 0.05$, **$P < 0.01$, ***$P < 0.001$ and ****$P < 0.0001$) were determined by non-parametric one-tailed Mann-Whitney U test or parametric one-tailed t-test depending on normality checks or two-way ANOVA with Sidak's multiple comparisons test for comparisons involving more than two groups. Outliers were detected using ROUT test (Q = 1 %). S = Start, F = Familiar, N = Novel.

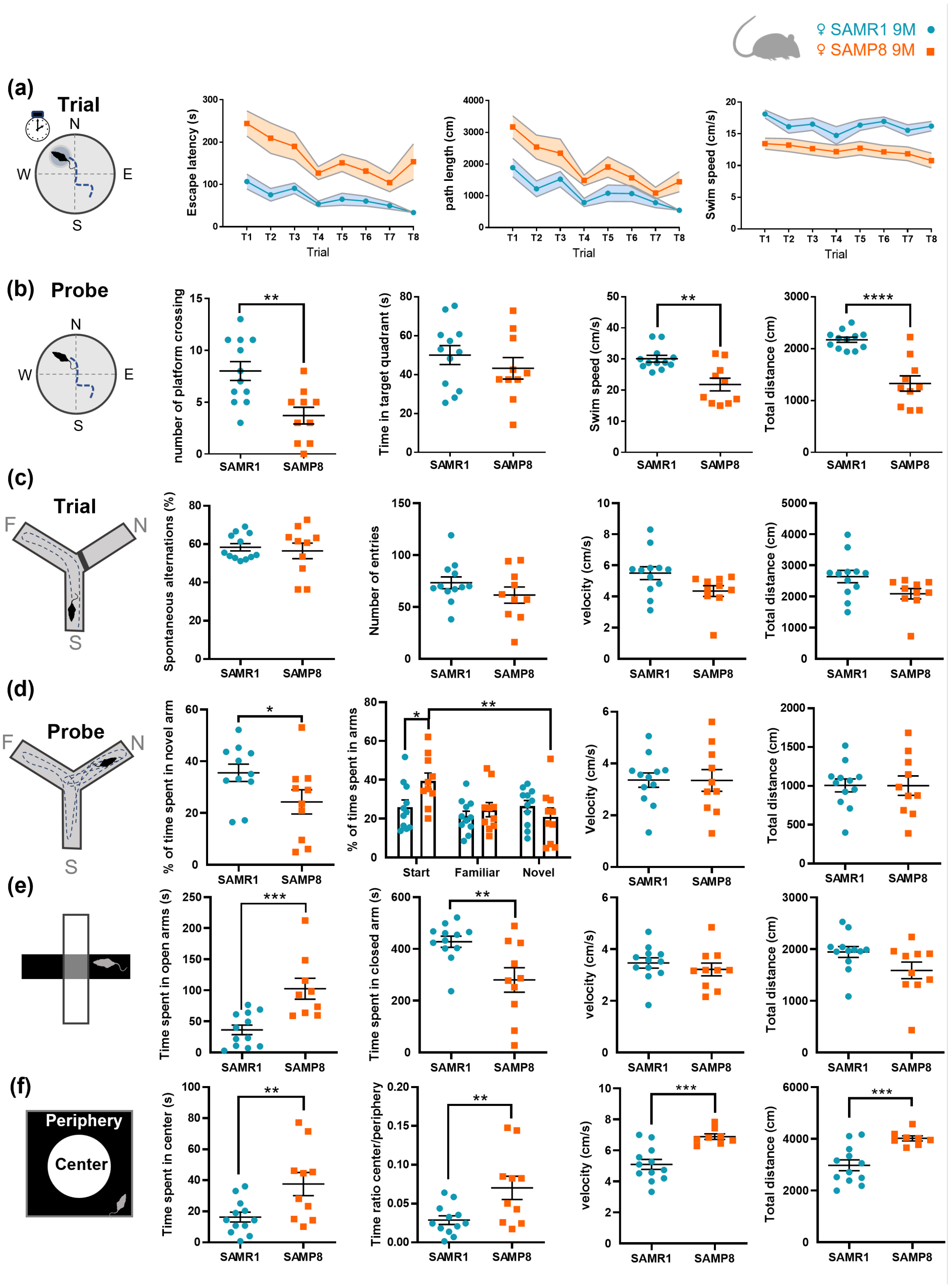
♀ SAMR1 9M
♀ SAMP8 9M
(a)
Trial
N
W
E
S
Escape latency (s)
path length (cm)
Swim speed (cm/s)
Trial
(b)
Probe
number of platform crossing
Time in target quadrant (s)
Swim speed (cm/s)
Total distance (cm)
SAMR1
SAMP8
(c)
Trial
F
N
S
Spontaneous alternations (%)
Number of entries
velocity (cm/s)
Total distance (cm)
(d)
Probe
% of time spent in novel arm
% of time spent in arms
Start
Familiar
Novel
Velocity (cm/s)
Total distance (cm)
(e)
Time spent in open arms (s)
Time spent in closed arm (s)
velocity (cm/s)
Total distance (cm)
(f)
Periphery
Center
Time spent in center (s)
Time ratio center/periphery
velocity (cm/s)
Total distance (cm)

**Supplementary Figure 8. Characterization of isolated CD11b+ cells from SAMR1 and SAMP8 mice.** (**a**) UMAP plot of CD11b+ cells depicting the different cell types of all treated groups (51182 cells). (**b**) Dot plot displaying the representative protein markers of the different cluster. (**c**) Dot plot depicting the top 10 differentially expressed genes of each cluster. (**d**) Feature Plot representing specific markers of each cell type.

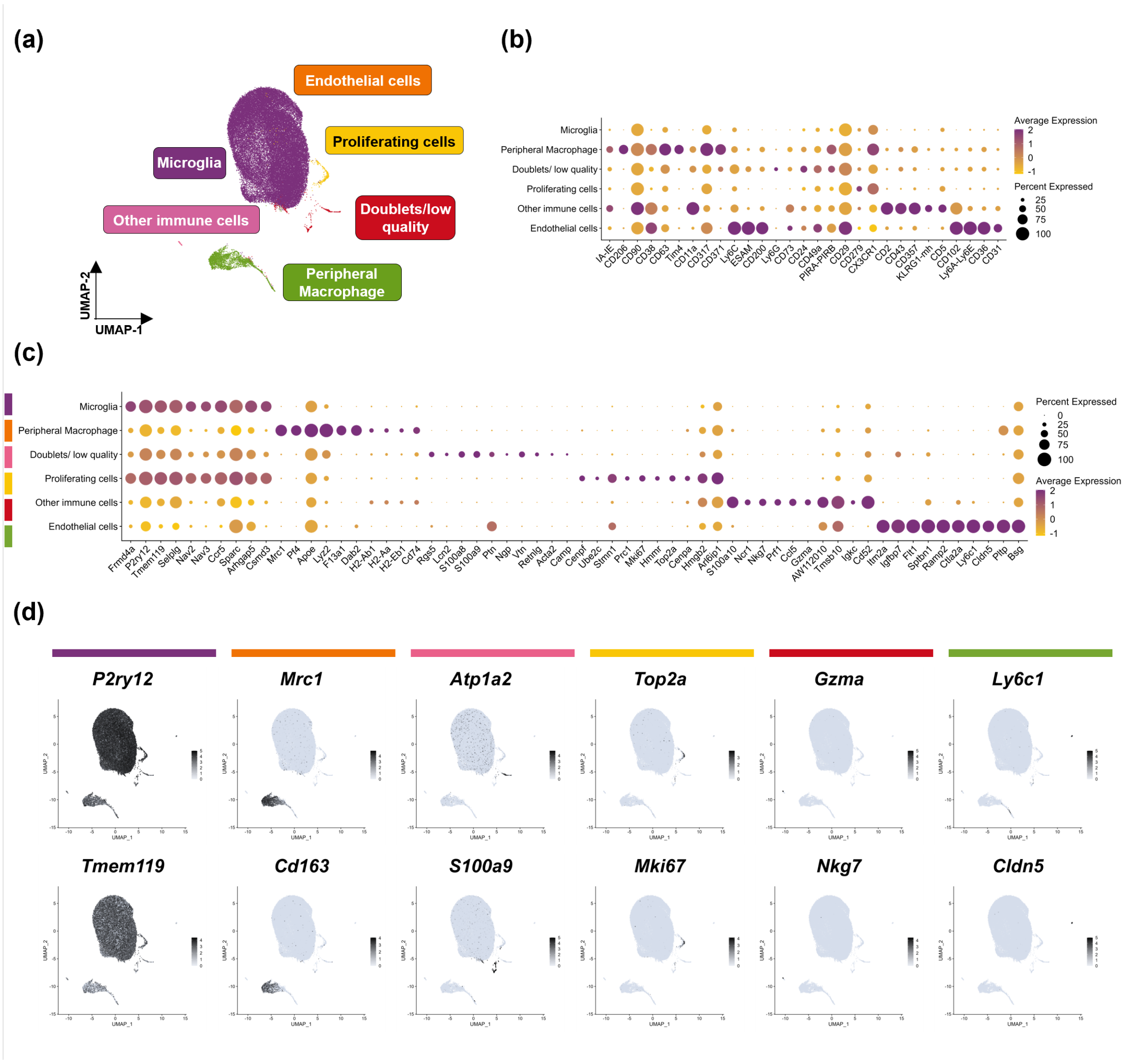

**Supplementary Figure 9. Differential expression between SAMR1 and SAMP8 at different timepoint.** (**a**-**c**) Volcano plot representing the differentially expressed genes in microglia between SAMP8 and SAMR1 mice groups at 2-months-old (**a**), 5-months-old (**b**), 9-months-old (**c**). (**d**) Volcano plot depicting the differentially expressed genes in microglia between 9-months SAMP8 (9M-SAMP8) and 2-months SAMP8 mice (2M-SAMP8).

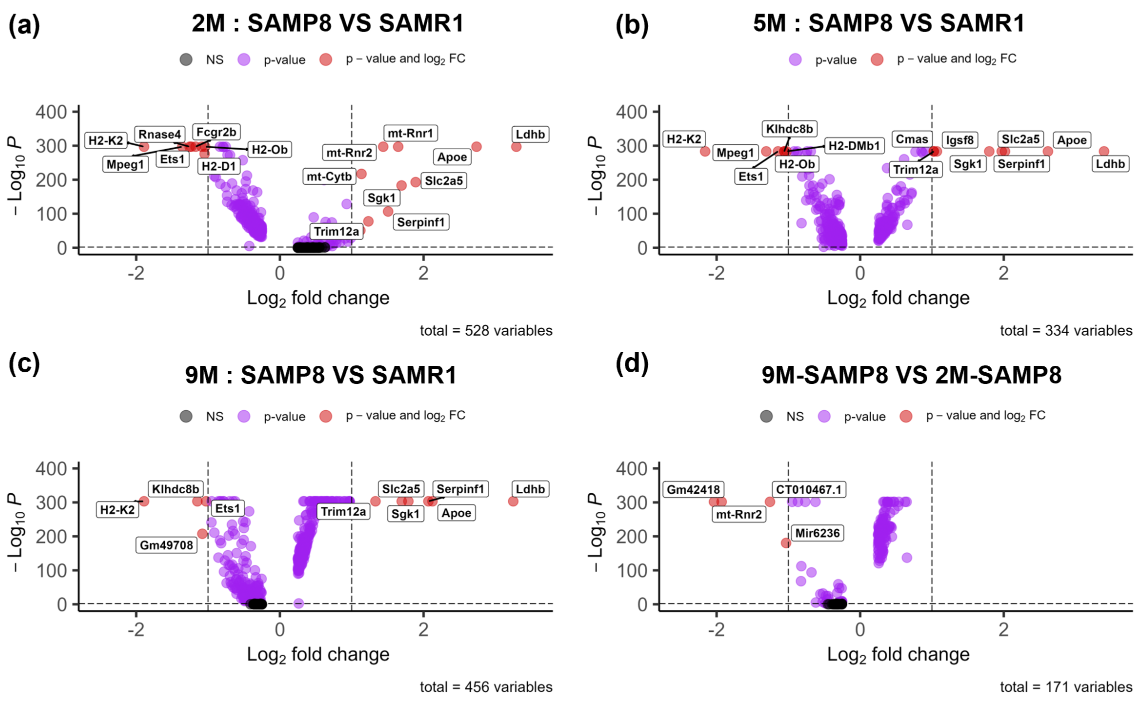

### Supplementary Tables

**Supplementary Table 1 Overview of mouse models, injections and treatments schemes used in this study.** References to the mouse models are provided in the material and method section. *M* = months.

| Mouse model | | Treatment | number of mice (n) | Strain | Sex | Endpoint | |
|---|---|---|---|---|---|---|---|
| Model | IC Injection at 3M | | | | | Age (M) | Experiment |
| CX3CR1$^{+/GFP}$ | | modified AIN76A + PBS (7 days) | 3 | C55BL/6J | Male | 2M | Flow cytometry (CX3CR1-GFP) |
| CX3CR1$^{+/GFP}$ | | modified AIN76A + PBS (14 days) | 3 | C55BL/6J | Male | 2M | Flow cytometry (CX3CR1-GFP) |
| CX3CR1$^{+/GFP}$ | | modified AIN76A + PBS (21 days) | 4 | C55BL/6J | Male | 2M | Flow cytometry (CX3CR1-GFP) |
| CX3CR1$^{+/GFP}$ | | modified AIN76A + PBS (28 days) | 5 | C55BL/6J | Male | 2M | Flow cytometry (CX3CR1-GFP) + Confocal Microscopy (IBA1, NEUN, SOX10, SOX9) on sections |
| CX3CR1$^{+/GFP}$ | | modified AIN76A + PLX5622 (7 days) | 2 | C55BL/6J | Male | 2M | Flow cytometry (CX3CR1-GFP) |
| CX3CR1$^{+/GFP}$ | | modified AIN76A + PLX5622 (14 days) | 5 | C55BL/6J | Male | 2M | Flow cytometry (CX3CR1-GFP) |
| CX3CR1$^{+/GFP}$ | | modified AIN76A + PLX5622 (21 days) | 5 | C55BL/6J | Male | 2M | Flow cytometry (CX3CR1-GFP) |
| CX3CR1$^{+/GFP}$ | | modified AIN76A + PLX5622 (28 days) | 5 | C55BL/6J | Male | 2M | Flow cytometry (CX3CR1-GFP) + Confocal Microscopy (IBA1, NEUN, SOX10, SOX9) on sections |
| CX3CR1$^{+/GFP}$ | | modified AIN76A + PLX3397 (7 days) | 3 | C55BL/6J | Male | 2M | Flow cytometry (CX3CR1-GFP) |
| CX3CR1$^{+/GFP}$ | | modified AIN76A + PLX3397 (14 days) | 4 | C55BL/6J | Male | 2M | Flow cytometry (CX3CR1-GFP) |
| CX3CR1$^{+/GFP}$ | | modified AIN76A + PLX3397 (21 days) | 5 | C55BL/6J | Male | 2M | Flow cytometry (CX3CR1-GFP) |
| CX3CR1$^{+/GFP}$ | | modified AIN76A + PLX3397 (28 days) | 4 | C55BL/6J | Male | 2M | Flow cytometry (CX3CR1-GFP) + Confocal Microscopy (IBA1, NEUN, SOX10, SOX9) on sections |
| C57Bl6/J | | | 9 | C55BL/6J | Female | 6M | qPCR (*Dusp1*, *Egr1*, *Fos*, *Fosb*, *Nfr4a1*, *Tnfa*, *Il1b*) |
| FVB | +PBS | | 4 | FVB | Male | 3M | CITE-Seq |
| FVB | +PBS | | 4 | FVB | Male | 4M | CITE-Seq |
| FVB | +PBS | | 4 | FVB | Male | 6M | CITE-Seq |

| | | | | | | | |
|---|---|---|---|---|---|---|---|
| FVB | +PBS | modified AIN76A + PBS (14 days pre- + 3 months post-injection) | 3 | FVB | Male | 6M | 3D microscopy (AT8) on hemibrains |
| FVB | +PBS | modified AIN76A + PBS (14 days pre- + 3 months post-injection) | 6 | FVB | Male | 6M | Confocal Microscopy (IBA1, S100A8/MRP8, CD63) on sections |
| FVB | +PBS | modified AIN76A + PLX3397 (14 days pre- + 3 months post-injection) | 3 | FVB | Male | 6M | Confocal Microscopy (IBA1) on sections |
| FVB | +PBS | | 4 | FVB | Male | 4M | Confocal Microscopy (IBA1, CD63) on sections |
| P301L | +PBS | | 4 | FVB | Male | 3M | CITE-Seq |
| P301L | +PBS | | 4 | FVB | Male | 4M | CITE-Seq |
| P301L | +PBS | | 4 | FVB | Male | 6M | CITE-Seq |
| P301L | +PBS | modified AIN76A + PBS (14 days pre- + 3 months post-injection) | 3 | FVB | Male | 6M | 3D microscopy (AT8) on hemibrains |
| P301L | +PBS | modified AIN76A + PBS (14 days pre- + 3 months post-injection) | 6 | FVB | Male | 6M | Confocal Microscopy (IBA1, S100A8/MRP8, CD63) on sections |
| P301L | +PBS | modified AIN76A + PLX3397 (14 days pre- + 3 months post-injection) | 3 | FVB | Male | 6M | Confocal Microscopy (IBA1) on sections |
| P301L | +PBS | | | FVB | Male | 4M | Confocal Microscopy (IBA1, CD63) on sections |
| P301L | +K18 | | 4 | FVB | Male | 3M | CITE-Seq |
| P301L | +K18 | | 4 | FVB | Male | 4M | CITE-Seq |
| P301L | +K18 | | 4 | FVB | Male | 6M | CITE-Seq |
| P301L | +K18 | modified AIN76A + PBS (14 days pre- + 3 months post-injection) | 7 | FVB | Male | 6M | 3D microscopy (AT8) on hemibrains |
| P301L | +K18 | modified AIN76A + PLX3397 (14 days pre- + 3 months post-injection) | 7 | FVB | Male | 6M | 3D microscopy (AT8) on hemibrains |
| P301L | +K18 | modified AIN76A + PBS (14 days pre- + 3 months post-injection) | 9 | FVB | Male | 6M | Confocal Microscopy (IBA1, S100A8/MRP8, CD63) on sections |
| P301L | +K18 | modified AIN76A + PLX3397 (14 days pre- + 3 months post-injection) | 3 | FVB | Male | 6M | Confocal Microscopy (IBA1) on sections |
| P301L | +K18 | | 4 | FVB | Male | 4M | Confocal Microscopy (IBA1, CD63) on sections |

| | | | | | | | |
|---|---|---|---|---|---|---|---|
| SAMR1 | | | 4 | AKR/J | Female | 2M | CITE-Seq |
| SAMR1 | | | 4 | AKR/J | Female | 5M | CITE-Seq |
| SAMR1 | | | 4 | AKR/J | Female | 9M | CITE-Seq |
| SAMR1 | | | 12 | AKR/J | Female | 9M | Behavioural tests (n =12) + Confocal Microscopy (S100A8) on sections (n=6) |
| SAMR1 | | | 3 | AKR/J | Male | 9M | Confocal Microscopy (pFTAA) on sections |
| SAMR1 | | | 3 | AKR/J | Male | 9M | Immunochemistry (PAS, 4G8, DAB staining) on sections |
| SAMP8 | | | 4 | AKR/J | Female | 2M | CITE-Seq |
| SAMP8 | | | 4 | AKR/J | Female | 5M | CITE-Seq |
| SAMP8 | | | 4 | AKR/J | Female | 9M | CITE-Seq |
| SAMP8 | | | 10 | AKR/J | Female | 9M | Behavioural tests (n =10) + Confocal Microscopy (S100A8) on sections (n=5) |
| SAMP8 | | | 3 | AKR/J | Male | 9M | Confocal Microscopy (pFTAA) on sections |
| SAMP8 | | | 3 | AKR/J | Male | 9M | Immunochemistry (PAS, 4G8, DAB staining) on sections |

**Supplementary Table 2 List of antibodies used for CITE-Seq.**

| TotalSeq | Barcode | Description | Clone | Isotype |
|---|---|---|---|---|
| A | 0001 | CD4 | RM4-5 | Rat IgG2a, κ |
| A | 0002 | CD8a | 53-6.7 | Rat IgG2a, κ |
| A | 0003 | CD366 | RMT3-23 | Rat IgG2a, κ |
| A | 0004 | CD279 | RMP1-30 | Rat IgG2b, κ |
| A | 0012 | CD117 | 2B8 | Rat IgG2b, κ |
| A | 0013 | Ly-6C | HK1.4 | Rat IgG2c, κ |
| A | 0014 | CD11b | M1/70 | Rat IgG2b, κ |
| A | 0015 | Ly-6G | 1A8 | Rat IgG2a, κ |
| A | 0070 | CD49f | GoH3 | Rat IgG2a, κ |
| A | 0074 | CD54 | YN1/1.7.4 | Rat IgG2b, κ |
| A | 0075 | CD90.2 | 30-H12 | Rat IgG2b, κ |
| A | 0076 | CD15 | MC-480 | Mouse IgM, κ |
| A | 0077 | CD73 | TY/11.8 | Rat IgG1, κ |
| A | 0078 | CD49d | R1-2 | Rat IgG2b, κ |
| A | 0079 | CD200 | OX-90 | Rat IgG2a, κ |
| A | 0090 | IgG1, κ Isotype Ctrl | MOPC-21 | Mouse (BALB/c) IgG1, Œ∫ |
| A | 0091 | IgG2a, κ Isotype Ctrl | MOPC-173 | Mouse IgG2a, Œ∫ |
| A | 0092 | IgG2b, κ Isotype Ctrl | MPC-11 | Mouse IgG2b, Œ∫ |
| A | 0093 | CD19 | 6D5 | Rat IgG2a, κ |
| A | 0095 | IgG2b, κ Isotype Ctrl | RTK4530 | Rat IgG2b, Œ∫ |
| A | 0097 | CD25 | PC61 | Rat IgG1, λ |
| A | 0098 | CD135 | A2F10 | Rat IgG2a, κ |
| A | 0103 | CD45R/B220 | RA3-6B2 | Rat IgG2a, κ |
| A | 0104 | CD102 | 3C4 (MIC2/4) | Rat IgG2a, κ |
| A | 0105 | CD115 | AFS98 | Rat IgG2a, κ |
| A | 0106 | CD11c | N418 | Armenian Hamster IgG |

| | | | | |
|---|---|---|---|---|
| A | 0107 | CD21,CD35 | 7E9 | Rat IgG2a, κ |
| A | 0108 | CD23 | B3B4 | Rat IgG2a, κ |
| A | 0109 | CD16/32 | 93 | Rat IgG2a, λ |
| A | 0110 | CD43 | S11 | Rat IgG2b |
| A | 0111 | CD5 | 53-7.3 | Rat IgG2a, κ |
| A | 0112 | CD62L | MEL-14 | Rat IgG2a, κ |
| A | 0113 | CD93 | AA4.1 | Rat IgG2b, κ |
| A | 0114 | F4/80 | BM8 | Rat IgG2a, κ |
| A | 0115 | FcεRIα | MAR-1 | Armenian Hamster IgG |
| A | 0117 | I-A/I-E | M5/114.15.2 | Rat IgG2b, κ |
| A | 0118 | NK-1.1 | PK136 | Mouse IgG2a, κ |
| A | 0119 | Siglec H | 551 | Rat IgG1, κ |
| A | 0120 | TCR β chain | H57-597 | Armenian Hamster IgG |
| A | 0121 | TCR γ/δ | GL3 | Armenian Hamster IgG |
| A | 0122 | TER-119 | TER-119 | Rat IgG2b, κ |
| A | 0130 | Ly-6A/E | D7 | Rat IgG2a, κ |
| A | 0134 | CD146 | P1H12 | Mouse IgG1, κ |
| A | 0171 | CD278 | C398.4A | Armenian Hamster IgG |
| A | 0173 | CD206 | C068C2 | Rat IgG2a, κ |
| A | 0182 | CD3 | 17A2 | Rat IgG2b, κ |
| A | 0184 | CD335 | 29A1.4 | Rat IgG2a, κ |
| A | 0190 | CD274 | MIH6 | Rat IgG2a, κ |
| A | 0191 | CD27 | LG.3A10 | Armenian Hamster IgG |
| A | 0192 | CD20 | SA275A11 | Rat IgG2b, κ |
| A | 0193 | CD357 | DTA-1 | Rat IgG2b, λ |
| A | 0194 | CD137 | 17B5 | Syrian Hamster IgG |
| A | 0195 | CD134 | OX-86 | Rat IgG1, κ |
| A | 0197 | CD69 | H1.2F3 | Armenian Hamster IgG |

| | | | | |
|---|---|---|---|---|
| A | 0198 | CD127 | A7R34 | Rat IgG2a, κ |
| A | 0200 | CD86 | GL-1 | Rat IgG2a, κ |
| A | 0201 | CD103 | 2E7 | Armenian Hamster IgG |
| A | 0202 | CD64 | X54-5/7.1 | Mouse IgG1, κ |
| A | 0203 | CD150 | TC15-12F12.2 | Rat IgG2a, λ |
| A | 0209 | TCR Vγ1.1 | 2.11 | Armenian Hamster IgG |
| A | 0210 | TCR Vγ3 | 536 | Syrian Hamster IgG |
| A | 0211 | TCR Vγ2 | UC3-10A6 | Armenian Hamster IgG |
| A | 0212 | CD24 | M1/69 | Rat IgG2b, κ |
| A | 0214 | Integrin β7 | FIB504 | Rat IgG2a, κ |
| A | 0222 | ERK1 | W15133A | Rat IgG2a, κ |
| A | 0223 | RORg | 2F7-2 | Mouse IgG2a, κ |
| A | 0225 | CD196 | 29-2L17 | Armenian Hamster IgG |
| A | 0226 | CD106 | 429 (MVCAM.A) | Rat IgG2a, κ |
| A | 0229 | CD62P | RMP-1 | Mouse IgG2a, κ |
| A | 0230 | CD8b | YTS156.7.7 | Rat IgG2b, κ |
| A | 0232 | MAdCAM-1 | MECA-367 | Rat IgG2a, κ |
| A | 0235 | TCR Vβ8.1,8.2 | KJ16-133.18 | Rat IgG2a, κ |
| A | 0236 | IgG1, κ Isotype Ctrl | RTK2071 | Rat IgG1, Œ∫ |
| A | 0237 | IgG1, λ Isotype Ctrl | G0114F7 | Rat IgG1, Œª |
| A | 0238 | IgG2a, κ Isotype Ctrl | RTK2758 | Rat IgG2a, Œ∫ |
| A | 0239 | IgG2a | #N/A | #N/A |
| A | 0240 | Rat IgG2c, κ Isotype Ctrl | RTK4174 | Rat IgG2c, Œ∫ |
| A | 0241 | IgG Isotype Ctrl | HTK888 | Armenian Hamster IgG |
| A | 0249 | IRF4 | IRF4.3E4 | Rat IgG1, κ |
| A | 0250 | KLRG1 | 2F1/KLRG1 | Syrian Hamster IgG |
| A | 0354 | TCR Vβ5.1, 5.2 | MR9-4 | Mouse IgG1, κ |
| A | 0376 | CD195 | HM-CCR5 | Armenian Hamster IgG |

| | | | | |
|---|---|---|---|---|
| A | 0378 | CD223 | C9B7W | Rat IgG1, κ |
| A | 0379 | CD62E | RME-1/CD62E | Mouse IgG1, κ |
| A | 0380 | CD90/CD90.1 | OX-7 | #N/A |
| A | 0381 | Panendothelial Cell Antigen | MECA-32 | Rat IgG2a, κ |
| A | 0388 | CD152 | UC10-4B9 | Armenian Hamster IgG |
| A | 0415 | P2RY12 | S16007D | Rat IgG2b, κ |
| A | 0416 | CD300LG | ZAQ5 | Rat IgG2a, κ |
| A | 0417 | CD163 | S15049I | Rat IgG2a, κ |
| A | 0421 | CD49b | HMα2 | Armenian Hamster IgG |
| A | 0422 | CD172a (SIRPα) | P84 | Rat IgG1, κ |
| A | 0424 | CD14 | Sa14-2 | Rat IgG2a, κ |
| A | 0426 | CD192 (CCR2) | SA203G11 | Rat IgG2b, κ |
| A | 0429 | CD48 | HM48-1 | Armenian Hamster IgG |
| A | 0434 | ReceptorD4 | #N/A | #N/A |
| A | 0435 | GABRB3 | #N/A | #N/A |
| A | 0439 | CD201 | RCR-16 | Rat IgG2a, κ |
| A | 0440 | CD169 | 3D6.112 | Rat IgG2a, κ |
| A | 0441 | CD71 | RI7217 | Rat IgG2a, κ |
| A | 0442 | Notch 1 | HMN1-12 | Armenian Hamster IgG |
| A | 0443 | CD41 | MWReg30 | Rat IgG1, κ |
| A | 0444 | CD184 | L276F12 | Rat IgG2b, κ |
| A | 0448 | CD204 | 1F8C33 | Rat IgG2a |
| A | 0449 | CD326 | G8.8 | Rat IgG2a, κ |
| A | 0450 | IgM | RMM-1 | Rat IgG2a, κ |
| A | 0551 | CD301a | LOM-8.7 | Rat IgG2a, κ |
| A | 0554 | CD309 | 89B3A5 | Rat IgG2a, κ |
| A | 0555 | CD36 | HM36 | Armenian Hamster IgG |
| A | 0556 | CD370 | 7H11 | Rat IgG1, κ |

| | | | | |
|---|---|---|---|---|
| A | 0557 | CD38 | 90 | Rat IgG2a, κ |
| A | 0558 | CD55 | RIKO-3 | Armenian Hamster IgG |
| A | 0559 | CD63 | NVG-2 | Rat IgG2a, κ |
| A | 0560 | CD68 | FA-11 | Rat IgG2a |
| A | 0561 | CD79b | HM79-12 | Armenian Hamster IgG |
| A | 0562 | CD83 | Michel-19 | Rat IgG1, κ |
| A | 0563 | CX3CR1 | SA011F11 | Mouse IgG2a, κ |
| A | 0564 | Folate Receptor β | 10/FR2 | Rat IgG2a, κ |
| A | 0565 | MERTK | 2B10C42 | Rat IgG2a, κ |
| A | 0566 | CD301b | URA-1 | Rat IgG2a, λ |
| A | 0567 | Tim-4 | RMT4-54 | Rat IgG2a, κ |
| A | 0568 | XCR1 | ZET | Mouse IgG2b, κ |
| A | 0570 | CD29 | HMβ1-1 | Armenian Hamster IgG |
| A | 0571 | IgD | 11-26c.2a | Rat IgG2a, κ |
| A | 0573 | CD140a | APA5 | Rat IgG2a, κ |
| A | 0595 | CD11a | M17/4 | Rat IgG2a, κ |
| A | 0596 | ESAM | 1G8/ESAM | Rat IgG2a, κ |
| A | 0807 | CD200R | OX-110 | Rat IgG2a, κ |
| A | 0808 | CD193 | J073E5 | Rat IgG2a, κ |
| A | 0809 | CD200R3 | Ba13 | Rat IgG2a, κ |
| A | 0810 | CD138 | 281-2 | Rat IgG2a, κ |
| A | 0811 | CD317 | 927 | Rat IgG2b, κ |
| A | 0812 | CD105 | MJ7/18 | Rat IgG2a, κ |
| A | 0813 | CD9 | MZ3 | Rat IgG2a, κ |
| A | 0824 | P2X7R | 1F11 | Rat IgG2b, κ |
| A | 0825 | CD371 | 5D3/CLEC12A | Rat IgG2a, κ |
| A | 0827 | CD22 | OX-97 | Rat IgG1, κ |
| A | 0834 | CD39 | Duha59 | Rat IgG2a, κ |

| | | | | |
|---|---|---|---|---|
| A | 0835 | CD314 | CX5 | Rat IgG1, κ |
| A | 0836 | DR3 | 4C12 | Armenian Hamster IgG |
| A | 0837 | IL-33Rα | DIH9 | Rat IgG2a, κ |
| A | 0846 | CD185 | L138D7 | Rat IgG2b, κ |
| A | 0848 | TIGIT | 1G9 | Mouse IgG1, κ |
| A | 0849 | CD80 | 16-10A1 | Armenian Hamster IgG |
| A | 0851 | CD1d | 1B1 | Rat IgG2b, κ |
| A | 0852 | CD226 | 10E5 | Rat IgG2b, κ |
| A | 0876 | CD300c/d | TX52 | Rat IgG2b, κ |
| A | 0877 | JAML | 4E10 | Armenian Hamster IgG |
| A | 0881 | CD272 | 6A6 | Armenian Hamster IgG |
| A | 0882 | PIR-A/B | 6C1 | Rat IgG1, κ |
| A | 0885 | CD270 | HMHV-1B18 | Armenian Hamster IgG |
| A | 0890 | CD137L | TKS-1 | Rat IgG2a, κ |
| A | 0891 | ENPP1 | YE1/19.1 | Rat IgG2b, κ |
| A | 0892 | CD2 | RM2-5 | Rat IgG2b, λ |
| A | 0905 | CD107a | 1D4B | Rat IgG2a, κ |
| A | 0916 | CD124 | I015F8 | Rat IgG2b, κ |
| A | 0917 | CD95 (Fas) | SA367H8 | Mouse IgG1, κ |

**Supplementary Table 3 Summary of sequencing parameters and cell counts after quality filtration.** *g/c* = gene per cell, *M* = months, *ncells* = number of cells; *Seq* = sequencing; *W* = weeks

| | | Group | FVB +PBS | FVB +PBS | FVB +PBS | P301L +PBS | P301L +PBS | P301L +PBS | P301L +K18 | P301L +PBS | P301L +K18 | SAMR1 | SAMR1 | SAMR1 | SAMP8 | SAMP8 | SAMP8 |
|---|---|---|---|---|---|---|---|---|---|---|---|---|---|---|---|---|---|
| | | Age | 1W | 1M | 3M | 1W | 1M | 3M | 1W | 1M | 3M | 2M | 5M | 9M | 2M | 5M | 9M |
| 1st seq. | | Cellranger QC nCells | 6,552 | 8,267 | 9,742 | 5,309 | 11,157 | 10,760 | 883 | 14,217 | 17,982 | 3,240 | 7,231 | 16,199 | 9,528 | 10,463 | 20,679 |
| | | mean Read/cells | 16,736 | 9,729 | 10,949 | 21,925 | 9,825 | 15,294 | 86,433 | 7,691 | 12,678 | 23,095 | 37,531 | 10,964 | 7,757 | 28,649 | 9,765 |
| | | Median g/c | 558 | 1,232 | 1,257 | 584 | 1,244 | 1,518 | 428 | 1,009 | 1,338 | 1,526 | 2,002 | 678 | 718 | 1,987 | 938 |
| 2nd seq. | | Cellranger QC nCells | | 10,164 | 10,556 | | 13,310 | 11,346 | | 20,456 | 19,838 | | | 18,257 | 19,454 | | 24,852 |
| | | mean Read/cells | | 24,544 | 28,034 | | 24,565 | 27,131 | | 22,635 | 27,510 | | | 27,789 | 14,438 | | 28,000 |
| | | Median g/c | | 1,930 | 1,995 | | 1,987 | 1,985 | | 1,794 | 1,931 | | | 1,014 | 810 | | 1,274 |
| | | Total integrated object (microglia) | 3,295 | 5,158 | 6,293 | 2,930 | 7,229 | 6,825 | | 8,724 | 10,709 | 1,491 | 3,800 | 11,334 | 11,717 | 5,799 | 13,059 |
| | | Animal 1 | 410 | 1,359 | 1,065 | 1,193 | 1,825 | 1,741 | | 1,833 | 2,768 | 493 | 940 | 3,095 | 3,386 | 1,221 | 2,116 |
| | | Animal 2 | 290 | 1,250 | 1,590 | 1,296 | 1,723 | 1,423 | | 2,322 | 2,256 | 343 | 931 | 3,130 | 2,529 | 2,046 | 4,010 |
| | | Animal 3 | 1,318 | 1,476 | 1,877 | 73 | 1,923 | 1,794 | | 2,466 | 2,894 | 231 | 969 | 3,160 | 3,048 | 1,444 | 2,631 |
| | | Animal 4 | 1,277 | 1,073 | 1,761 | 368 | 1,758 | 1,867 | | 2,103 | 2,791 | 424 | 960 | 1,949 | 2,754 | 1,088 | 4,302 |

**Supplementary Table 5 Overview of human samples.** F = Female, M = Male

| Group | ID number | Sex | Age (years) | Montine stage |
|---|---|---|---|---|
| Alzheimer's disease | 6159 | M | 59 | A3B3C3 |
| | 6474 | M | 57 | A3B3C3 |
| | 6044 | M | 57 | A3B3C3 |
| | 6190 | M | 67 | A3B3C3 |
| | 6427 | M | 62 | A3B3C3 |
| | 6373 | F | 53 | A3B3C3 |
| | 6374 | F | 57 | A3B3C3 |
| | 6406 | F | 51 | A3B3C3 |
| | 6217 | F | 63 | A3B3C3 |
| | 6126 | F | 65 | A3B3C3 |
| Tauopathy | 6225 | M | 53 | |
| | 6460 | M | 65 | |
| | 6280 | M | 70 | |
| | 6520 | M | 70 | |
| | 6113 | F | 51 | |
| | 6446 | F | 69 | |
| | 6452 | F | 67 | |
| | 6514 | F | 65 | |
| Healthy | 806 | M | 41 | |
| | 4985 | M | 47 | |
| | 4895 | M | 58 | |
| | 5875 | M | 59 | |
| | 5201 | M | 69 | |
| | 5633 | F | 44 | |
| | 4485 | F | 48 | |
| | 4480 | F | 59 | |
| | 3931 | F | 60 | |
| | 3686 | F | 69 | |

#### Supplementary Table 5 PCR primer sequences for RT-PCR.

| Gene symbol | Forward primer sequences (5'-3') | Reverse primer sequences (5'-3') |
|---|---|---|
| *Dusp1* | GTTGTTGGATTGTCGCTCCTT | TTGGGCACGATATGCTCCAG |
| *Egr1* | TCGGCTCCTTTCCTCACTCA | CTCATAGGGTTGTTCGCTCGG |
| *Fos* | CGGGTTTCAACGCCGACTA | TTGGCACTAGAGACGGACAGA |
| *Fosb* | TTTTCCCGGAGACTACGACTC | GTGATTGCGGTGACCGTTG |
| *Nrf4a1* | TTGAGTTCGGCAAGCCTACC | GTGTACCCGTCCATGAAGGTG |
| *Tnf* | CCTGTAGCCCACGTCGTAG | GGGAGTAGACAAGGTACAACCC |
| *Il1b* | TCTCGCAGCAGCACATCA | CACACACCAGCAGGTTAT |
| *Gapdh* | TGAAGGTCGGTGTGAACGG | CGTGAGTGGAGTCATACTGGAA |
| *Actb* | TGTCGAGTCGCGTCCACC | TCGTCATCCATGGCGAACTGG |
| *B2m* | ATTCACCCCCACTGAGACTG | TGCTATTTCTTTCTGCGTGC |
